# Pathogen-induced pH changes regulate the growth-defense balance of plants

**DOI:** 10.1101/550491

**Authors:** Christopher Kesten, Francisco M. Gámez-Arjona, Stefan Scholl, Alexandra Menna, Susanne Dora, Apolonio Ignacio Huerta, Hsin-Yao Huang, Nico Tintor, Toshinori Kinoshita, Martijn Rep, Melanie Krebs, Karin Schumacher, Clara Sánchez-Rodríguez

## Abstract

Environmental adaptation of organisms relies on fast perception and response to external signals, which lead to developmental changes. Plant cell growth is strongly dependent on cell wall remodeling. However, little is known about cell wall-related sensing of biotic stimuli and the downstream mechanisms that coordinate growth and defense responses. We generated genetically encoded pH sensors to determine absolute pH changes across the plasma membrane in response to biotic stress. A rapid apoplastic acidification by phosphorylation-based proton pump activation was followed by an acidification of the cortical side of the plasma membrane in response to the fungus *Fusarium oxysporum.* The proton chemical gradient modulation immediately reduced cellulose synthesis and cell growth and, furthermore, had a direct influence on the pathogenicity of the fungus. All these effects were dependent on the COMPANION OF CELLULOSE SYNTHASE proteins that are thus at the nexus of plant growth and defense. Hence, our discoveries show a remarkable connection between plant biomass production, immunity, and pH control, and advance our ability to investigate the plant growth-defense balance.

## Introduction

Developmental adaptation to the environment, such as growth and differentiation, is the driving force of evolution and key for the survival of organisms. Therefore, unraveling the mechanisms by which signal perception and response are coupled to growth at the cellular level is one of the central challenges in biology.

Plants adapt their growth to the environment through precisely controlled changes in cell expansion and division, which rely on the accurate remodeling of their cell walls (CWs). This strong, yet extensible, polysaccharide-based organelle corsets the plant, while connected with the plasma membrane (PM) and cytosol (Liu *et al*, 2015; Cosgrove, 2018). The CW, as the structure occupying most of the apoplastic space, is the first cellular compartment which encounters environmental and internal signals and is directly involved in perception, transduction and response to these stimuli (Wolf, 2017; Kesten *et al*, 2017). Cellulose is an essential constituent of plant CWs. It is composed of β-1,4-linked D-glucose chains that are synthesized and extruded to the apoplast by PM-localized cellulose synthase (CesA) complexes (CSCs; (McFarlane *et al*, 2014)). The catalytically-driven CSCs traverse the PM, a process that is guided by cortical microtubules (MTs) (Paredez *et al*, 2006; Watanabe *et al*, 2015). CSCs and cortical MTs are physically connected by at least two protein families, the CELLULOSE SYNTHASE-INTERACTIVE PROTEIN 1/POM2 (CSI1/POM2; Bringmann et al., 2012; Li et al., 2012) and COMPANION OF CELLULOSE SYNTHASE proteins (CCs) (Endler *et al*, 2015). Importantly, the CSCs and cortical MTs are interdependent, as alterations in one affects the activity of the other. This mutual support has been reported by several studies, especially under abiotic stress conditions such as drought and salt (Gutierrez *et al*, 2009; Nick, 2013; Endler *et al*, 2015; Wang *et al*, 2016). Recently, CCs were shown to be indispensable for the recovery of the cortical MT array and CSC activity under salt stress (Endler *et al*, 2015), and the microtubule interacting N-terminus of CC1 to be key for this function (Kesten *et al*, 2018).

The alteration of cellulose synthase activity in response to extra- and intracellular signals is vital for growth modification under dynamic environmental conditions. Changes in pH within distinct cell compartments, including the apoplast/CW, act as messengers to transfer such signals (Felle, 2001; Sze & Chanroj, 2018). Plants and fungi primarily regulate pH at across the PM via the activity of proton pumps, such as PM H^+^-ATPases. The active translocation of protons to the apoplast by these pumps provides the chemical and electrical potential needed for solute transport (Haruta *et al*, 2015). The activity of Arabidopsis H^+^-ATPases (AHAs), is essential for plant growth and development under different environmental conditions. Dynamic phosphorylation of specific AHA amino acid residues rapidly regulates proton pump activity in response to external and internal stimuli, such as abiotic and biotic stresses or hormones (Olsson *et al*, 1998; Gao *et al*, 2004; Jeworutzki *et al*, 2010; Stecker *et al*, 2014; Haruta *et al*, 2015; Falhof *et al*, 2016; Geilfus, 2017). For example, the activation of AHAs is known to require the phosphorylation of their penultimate C-terminal threonine residue (Inoue & Kinoshita, 2017). Moreover, rapid adjustment of the CW/apoplastic pH can influence CW loosening/stiffening, and thus the connection of the CW to the underlying cell (Fendrych *et al*, 2016; Barbez *et al*, 2017) by a mechanism which is not yet fully understood (Mangano *et al*, 2018). Interestingly, plant microbes have been shown to secrete various CW degrading enzymes depending on the apoplastic pH (pH_apo_) of their host (Li *et al*, 2012). In addition, the pH_apo_ plays an essential role in the infection process of some pathogens, such as the root vascular fungus *Fusarium oxysporum* (*Fo*) (Masachis *et al*, 2016; López-Díaz *et al*, 2018). *Fo* hyphae attach to the root surface, penetrate through natural openings, and then grow intercellularly towards xylem vessels where they can proliferate throughout the plant. Once in the xylem, *Fo* secretes a complex collection of proteins named Secreted In Xylem (SIX), which serve to evade plant defense responses (Pietro *et al*, 2003; Houterman *et al*, 2007). *Fo* can infect over a hundred crop species, with single strains infecting a narrow host-range. Among them, *F. oxysporum* 5176 (Fo5176) is one of the genetic strains that can infect Arabidopsis (Thatcher *et al*, 2012).

The regulation of cellulose synthesis is crucial for plant development and environmental adaptation. Yet the mechanisms underlying stress-induced changes in MT and CSC organization, their crosstalk with cellular signaling messengers, and broad cellular implications remain elusive, especially upon biotic stress (Kesten *et al*, 2017). This knowledge would be particularly relevant at the root level considering the essential role of this organ in nutrient uptake and response to abiotic and biotic stresses (Zamioudis *et al*, 2015; Feng *et al*, 2016; Smakowska *et al*, 2016; Morris *et al*, 2017). In this study, we use the Arabidopsis*-*Fo5176 pathosystem to characterize plant responses to microbes at a cellular level and how these influence plant adaptation to biotic stress.

## Results

### Rapid alteration of the cellulose synthesis machinery and cell elongation upon fungal contact is pH dependent

Plant CW reorganization in response to stress is essential for proper environmental adaptation. To determine the influence of biotic stress on the regulation of plant cellulose synthesis, we visualized CesAs and cortical MTs of *A. thaliana* roots exposed to Fo5176 hyphae. We aimed to mimic early contact events at the plant-microbe interface, and therefore used young hyphae obtained directly after spores germinated overnight. We observed a depletion of CesA foci at the PM (GFP-CesA3; (Desprez *et al*, 2007)) simultaneously to depolymerization of cortical MTs (mCherry-TUA5; (Gutierrez *et al*, 2009)) within five min of hypha contact, which were not observed in control roots (Fig. 1a-c; Supplementary Fig. 1a and b; Supplementary Movie 1). In addition to decreased CesA density, the speed of the remaining CesA foci at the PM was reduced in comparison to mock conditions (Supplementary Fig. 1b). In agreement with the observed molecular changes, roots pre-exposed to fungal hyphae for five min had a reduced root growth rate as compared to control plants (Fig. 1d and e). To exclude that the observed response was solely based on pressure applied by the hyphae to the roots, we prepared a Fo5176 elicitor mix containing solubilized molecules from a fungal culture (Baldrich *et al*, 2014). The fungal elicitor mix induced similar, rapid changes on the cellulose synthase machinery, as well as on cortical MTs, to those observed in response to alive hyphae, i.e. cortical MTs depolymerized and PM-localized CesA density and speed was reduced as compared to mock treated roots (Fig. 1f-h; Supplementary Fig. 1c; Supplementary Movie 2). Consequently, plants reacted with an immediate root growth reduction once the elicitors were added to the imaging media (Fig. 1i-j, Supplementary Movie 3), suggesting that a fungal elicitor induced the rapid changes in the CSCs-MT machinery and root growth.

**Figure 1:**
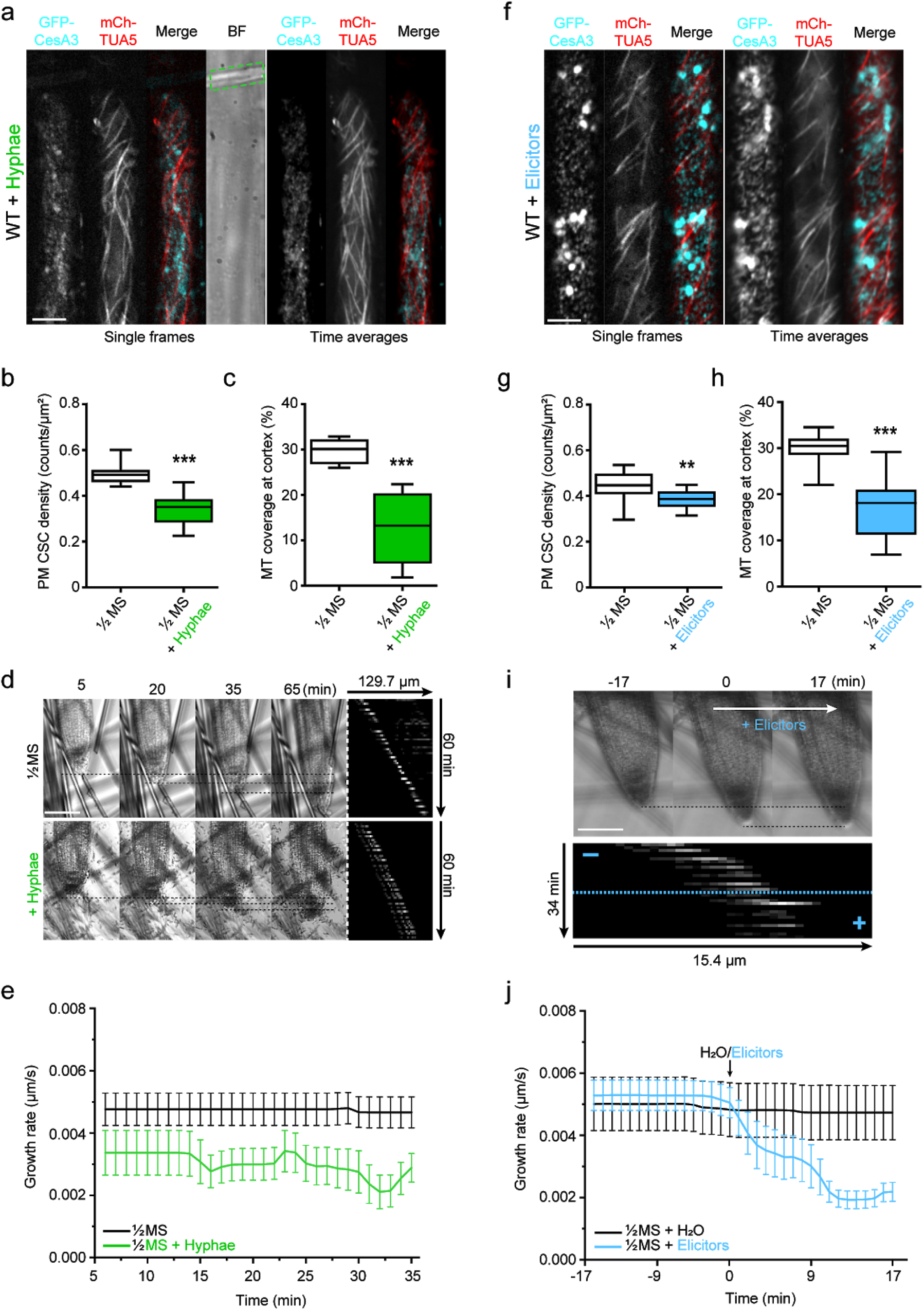
Fo5176 hyphae and elicitors cause immediate depletion of cellulose synthase complexes and cortical microtubules from the plasma membrane and cell cortex, simultaneously to growth rate reduction. **a** Representative image of a root epidermal cell from a 5 day-old wild-type (WT) GFP-CesA3 and mCh-TUA5 dual-labeled line upon 5 min of Fo5176 hyphae contact (left panels, single frame; right panels, time average projections). Fo5176 hypha is defined by a green dashed line in the brightfield (BF) channel. Scale bar = 5 μm. **b** Quantification of GFP-CesA3 density at the plasma membrane after Fo5176 hyphae contact as depicted in **a**. Box plots: center lines show the medians; box limits indicate the 25th and 75th percentiles; whiskers extend to the minimum and maximum. N ≥ 17 cells from 14 roots and 3 independent experiments. Welch’s unpaired *t*-test; *** p-value ≤ 0.001. **c** Quantification of microtubule density at the cell cortex, after Fo5176 hyphae contact as depicted in **a**. Box plots as described in **b**. N ≥ 8 cells from 8 roots and 3 independent experiments. Welch’s unpaired *t*-test; *** p-value ≤ 0.001. **d** Growth progression of roots grown in half MS or half MS + Fo5176 hyphae. Left panels: representative images of roots at different times after the corresponding treatment. Scale bar = 100 μm. Right panel: kymographs depicting growth of roots in the left panel. **e** Growth rate of roots in half MS or half MS + Fo5176 hyphae, analyzed from images as in **d**. Average growth rate in half MS: 0.0047 ± 0.0005 μm/s; average growth rate in half MS + Fo5176 hyphae: 0.0031 ± 0.0005 μm/s. Values are mean +/-SEM, N ≥ 11 seedlings from 3 independent experiments. Welch’s unpaired *t*-test; * p-value ≤ 0.05. **f** Representative image of a root epidermal cell from a WT GFP-CesA3 and mCh-TUA5 dual-labeled line upon 5 min elicitors treatment (left panels, single frame; right panels, time average projections). Scale bar = 5 μm. **g** Quantification of GFP-CesA3 density at the plasma membrane after elicitors treatment as depicted in **f**. Box plots as described in **b**. N ≥ 23 cells from 14 roots and 3 independent experiments. Welch’s unpaired *t*-test; ** p-value ≤ 0.01. **h** Quantification of microtubule density at the cell cortex, after elicitors treatment as depicted in **f**. Box plots as described in **b**. N ≥ 14 cells from 12 roots and 3 independent experiments. Welch’s unpaired *t*-test; *** p-value ≤ 0.001. **i** Growth progression of roots exposed to fungal elicitor mix. Upper panel: representative images of roots grown in half MS (−17 to 0 min) and after being exposed to the fungal elicitors added at 0 min. Scale bar = 100 μm. Lower panel: kymograph depicting growth of roots in the upper panel. **j** Growth rate of roots exposed to fungal elicitors, analyzed from images as in **a**. After 17 min of growth in half MS, H_2_O or elicitors were added and the growth rate was measured for an additional 17 min. Average growth rate before treatment (−17 to 0 min): H_2_O: 0.0050 ± 0.0009 μm/s; elicitor mix: 0.0053 ± 0.0005 μm/s. Average growth rate after treatment (0-17 min): H_2_O: 0.0048 ± 0.0009 μm/s; elicitor mix: 0.0028 ± 0.0004 μm/s. Values are mean +/-SEM, N ≥ 10 seedlings from 3 independent experiments. Welch’s unpaired *t*-test for roots before and after elicitors treatment; ** p-value ≤ 0.01.

Alteration of pH transients in distinct cell compartments is a fast cellular signalling event reported in response to stress (Behera *et al*, 2018). Therefore we investigated the influence of pH on the observed Fo5176 effect on roots. Reduction of CesA density and speed as well as cortical MT depolymerization were completely reversible by buffering the imaging media with 5 mM 2-(N-morpholino)ethanesulfonic acid-KOH (MES) (Fig. 2a-c; Supplementary Fig. 2a-c; Supplementary Movie S4), a common buffering substance in plant media (Good *et al*, 1966; Bugbee & Salisbury, 1985; Stahl *et al*, 1999). This implies the direct involvement of pH changes in the observed responses of plant roots to Fo5176.

**Figure 2:**
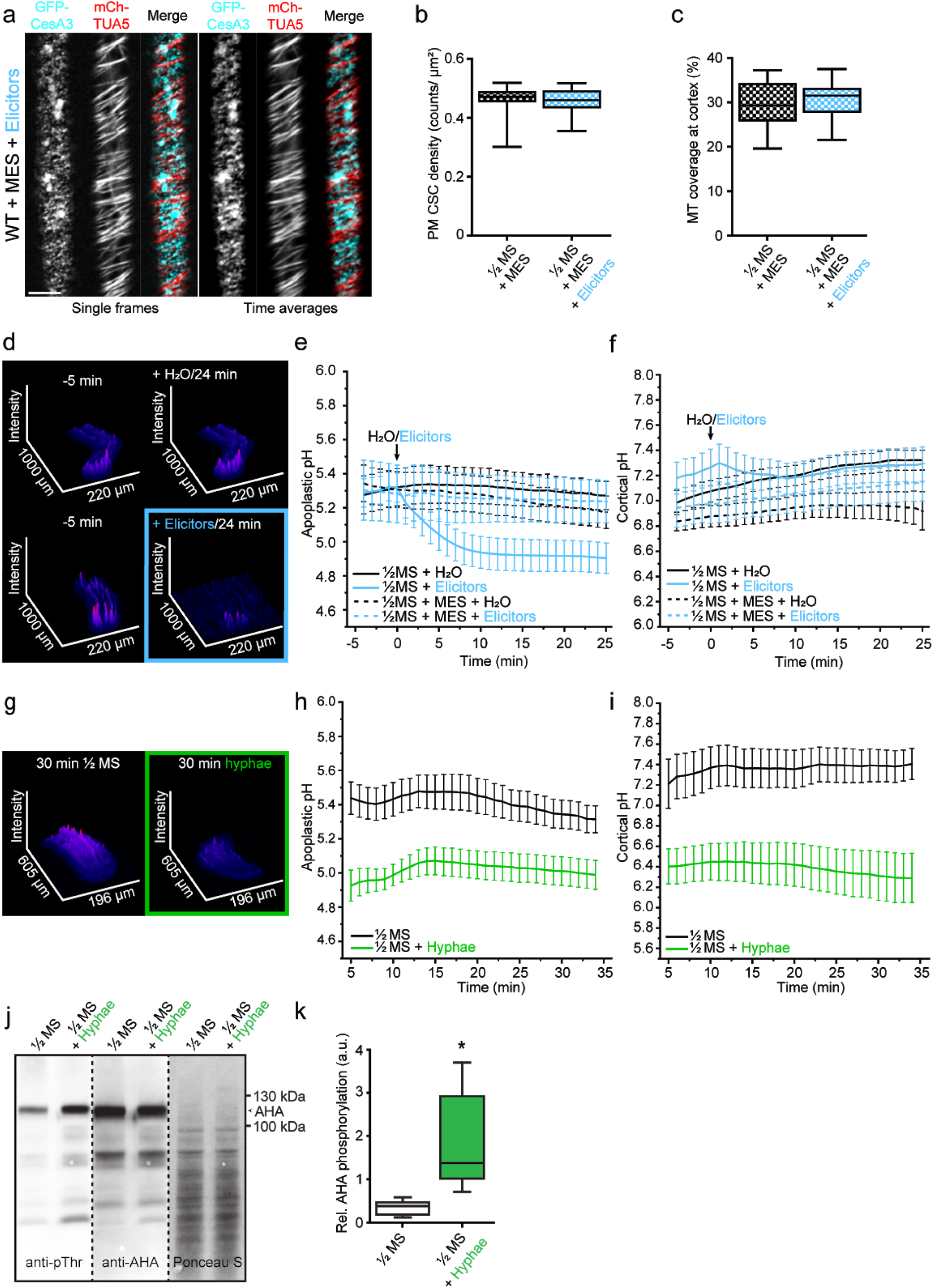
Fo5176-induced CSC-MT alterations and immediate acidification of the PM interface can be buffered with MES and are dependent on upregulation of AHA activity. **a** Representative image of a root epidermal cell from a 5 day-old wild-type (WT) GFP-CesA3 and mCh-TUA5 dual-labeled line upon 5 min of elicitors treatment in half MS + 5 mM MES (left panels: single frame; right panels: time average projections). Scale bar = 5 μm. **b** Quantification of GFP-CesA3 density at the plasma membrane after elicitors treatment in half MS + 5 mM MES as depicted in **a**. Box plots: center lines show the medians; box limits indicate the 25th and 75th percentiles; whiskers extend to the minimum and maximum. N ≥ 17 cells from 10 roots and 3 independent experiments. **c** Quantification of microtubule density at the cell cortex after elicitors treatment in half MS + 5 mM MES as depicted in **a**. Box plots as described in **b**. N ≥ 14 cells from 10 roots and 3 independent experiments. **d** Representative surface plot of a WT root expressing the pH_apo_ sensor SYP122-pHusion grown in half MS (−5 to 0 min). At 0 min either H_2_O (upper panel) or an elicitor mix (lower panel) was added. Upon elicitors treatment, the signal intensity in the depicted 488 nm channel drastically decreases (highlighted with a blue square). **e** Apoplastic pH variation in WT roots expressing the pH_apo_ sensor SYP122-pHusion over time, either in half MS (as depicted in **d**) or half MS + 5 mM MES. Imaging started 5 min before either H_2_O or a fungal elicitor mix was added (0 min). Values are mean +/-SEM, N ≥ 15 seedlings from 3 independent experiments. RM two-way ANOVA on half MS + H_2_O vs half MS + elicitors: p ≤ 0.05 (treatment), p ≤ 0.001 (time), p ≤ 0.001 (treatment x time). Please refer to Fig. S2i for ΔpH analysis. **f** Cortical pH variation of WT roots expressing the pH_cortical_ sensor pHGFP-Lti6b over time, either in half MS or half MS + 5 mM MES. Imaging started 5 min before either H_2_O or a fungal elicitor mix was added (0 min). Values are mean +/-SEM, N = 16 seedlings from 3 independent experiments. Mixed effects model on half MS + H_2_O vs half MS + elicitors: p = 0.80 (treatment), p ≤ 0.001 (time), p ≤ 0.001 (treatment x time). Please refer to Fig. S2j for ΔpH analysis. **g** Representative surface plot of WT root expressing the pH_apo_ sensor SYP122-pHusion grown 30 min in half MS (left panel) or Fo5176 hyphae (right panel). The hyphae treated root shows drastically reduced signal intensity in the depicted 488 nm channel (highlighted with a green square). **h** Apoplastic pH variation of WT roots expressing the pH_apo_ sensor SYP122-pHusion over time, either in half MS or half MS + Fo5176 hyphae. Roots were exposed to hyphae for 5 min before imaging started. The pH was calculated from image series as in **g**. Values are mean +/-SEM, N ≥ 12 seedlings from 3 independent experiments. RM two-way ANOVA on half MS vs half MS + elicitors: p ≤ 0.01 (treatment), p ≤ 0.001 (time), p ≤ 0.001 (treatment x time). **i** Cortical pH variation of WT roots expressing the pH_cortical_ sensor pHGFP-Lti6b over time, either in half MS or half MS Fo5176 hyphae. Roots were exposed to hyphae for 5 min before imaging started. Values are mean +/-SEM, N ≥ 13 seedlings from 3 independent experiments. RM two-way ANOVA on half MS vs half MS + elicitors: p ≤ 0.01 (treatment), p = 0.30 (time), p ≤ 0.001 (treatment x time). **j** Western blots showing chemiluminescent signals of anti-pThr or anti-AHA incubated membranes loaded with Arabidopsis root samples treated for 8 min with either half MS or half MS + Fo5176 hyphae. The Ponceau S panel shows total protein content. The AHA band used for quantification is highlighted with an arrowhead. Dashed line separates different treatments of the same membrane. **k** Quantification of relative (Rel.) AHA phosphorylation status from western blots as shown in **j,** calculated by obtaining the signal intensity ratio of anti-pThr in respect to anti-AHA. Box plots as described in **b**. N=5 independent experiments. Welch’s unpaired *t*-test; * p-value ≤ 0.05.

### New sensors to measure the pH at both sides of the plant PM

To test the above hypothesis, we sought to measure the pH of plant roots at different developmental zones, from meristem to mature regions, and various cell layers in response to alive Fo5176 hyphae and elicitors. As previously reported, the apoplastic pH sensor apo-pHusion suffers from high background signal in the endoplasmic reticulum (ER), which compromises precise pH measurements in the apoplast (Gjetting *et al*, 2012; Martinière *et al*, 2018). Therefore, we generated a pH_apo_ probe with as low as possible intracellular background signal by fusing the ratiometric probe pHusion (Gjetting *et al*, 2012) to the C-terminus of the PM-localized SNARE (soluble N-ethyl-maleimide sensitive factor attachment protein receptor) protein Syntaxin of Plants 122 (SYP122; (Assaad *et al*, 2004; Uemura *et al*, 2004)). Indeed, microscopic analysis of stable transgenic lines revealed considerably less intracellular signal for pUB10::SYP122-pHusion than for apo-pHusion (Supplementary Fig. 2d). *In vivo* calibration (Supplementary Fig. 2e) and pH measurements in control epidermal/cortex cells of the root elongation zone indicated an average pH_apo_ over five min of 5.30 ± 0.45 (Fig. 2 d-e; Values are mean +/- SD, N = 17 seedlings from 3 independent experiments), which is in good agreement with previously reported values obtained with fluorescent dyes (Barbez *et al*, 2017) or surface electrode measurements in the root elongation zone of Arabidopsis (Staal *et al*, 2011). To measure pH on both sides of the PM, we targeted the cortical side by fusing pHGFP (Moseyko & Feldman, 2001) to the N-terminus of the PM localized low temperature induced protein 6b (Lti6b;(Cutler *et al*, 2000), also under control of pUB10. Counterstaining of cell walls with propidium iodide (PI) indicated PM localization of pHGFP-Lti6b in root cells (Supplementary Fig. 2f). *In vivo* calibration of the cortical pH (pH_cortical_) sensor pHGFP-Lti6b revealed an almost linear relationship of emission ratio and pH in the range from pH 5.6 to 8.0 (Supplementary Fig. 2g). Cortical pH measurements in control epidermal/cortex cells of the root elongation zone indicated an average pH_cortical_ over 5 min of 7.03 ± 0.36 (Fig. 2f; Values are mean +/-SD, N = 16 seedlings from 3 independent experiments), which is in agreement with previous reports (Moseyko & Feldman, 2001; Gao *et al*, 2004; Schulte *et al*, 2006). Cytoplasmic pH (pH_cyto_) measurements were approached by expressing free pHGFP in the cytosol (Moseyko & Feldman, 2001; Fendrych *et al*, 2014). In control epidermal/cortex cells of the root elongation zone we measured an average pH_cyto_ over 5 min of 6.62 ± 0.92 (Supplementary Fig. 2h; Values are mean +/-SD, N = 15 seedlings from 3 independent experiments), which is also in agreement with previous reports using fluorescent methods (Moseyko & Feldman, 2001; Gao *et al*, 2004; Schulte *et al*, 2006).

### Fungal-induced pH changes across the PM are generated by the activation of proton pumps

Upon elicitors treatment, the pH_apo_ dropped below 5.0 within five min (Fig. 2d-e, Supplementary Fig. 2i; Supplementary Movie S5), simultaneously with the observed reduction of root growth (Fig. 1i-j). It is important to note that the pH of the added elicitor mix ranged between 5.5 to 5.8. The cortical side of the PM also acidified (Fig. 2f, Supplementary Fig. 2j), but the response was delayed by one minute in comparison to the apoplastic side. In this first minute, the pH_cortical_ rose as an immediate response to the acidification of the apoplast (Fig. 2f; Supplementary Fig. 2j), pointing towards a translocation of protons from the cortical to the apoplastic side of the PM. We could not detect any apparent changes in the pH_cyto_ in response to fungal elicitors (Supplementary Fig. 2h), highlighting that the immediate pH response was constraint to the PM environment. When we buffered the media with MES before adding the elicitor mix, no significant change of pH could be observed on both sides of the PM (Fig. 2e and f; Supplementary Fig. 2i and j). This suggests that the depletion of CSCs and cortical MTs, as well as the reduction of root growth upon elicitors treatment, is linked to PM pH changes.

Next, we explored if the cellular pH fluctuations were also detectable upon fungal contact. Therefore, we imaged plants on an agarose cushion that was coated with live Fo5176 hyphae. Under these conditions,the pH_apo_ and pH_cortical_ were comparable to the elicitor mix experiments under mock conditions (5.44 ± 0.33 and 6.99 ± 0.73, respectively). In line with the elicitor mix treated roots, five min of contact with Fo5176 hyphae also induced a drastic acidification of both the apoplast and the cortical side of the PM (Fig. 2g-i). The fast apoplastic acidification in response to Fo5176 points towards an increase in plant PM proton pump activity upon hyphae and elicitor mix contact. We approached this hypothesis by western blotting with well-characterized antibodies against the catalytic domain of the AHAs and their upregulated form that is phosphorylated on the penultimate threonine residue (Hayashi *et al*, 2010). Indeed, western blotting revealed increased phosphorylation of AHAs upon Fo5176 hyphae contact (Fig 2j and k), supporting the observed pH_apo_ drop upon Fo5176 exposure (Fig 2g and h). These results suggest that a ∆pH change across the PM induced the detected CSC depletion, cortical MT depolymerization, and root growth inhibition upon hyphae and elicitors treatment. We next mimicked a ∆pH change across the PM by employing a buffer that adjusts the external and internal pH of cells by the partitioning effect of a weak acid and weak base exchange (Buffer A: 25 mM MES-BTP pH 5.4, 25 mM CH_3_COONH_4_) (Heiple & Taylor, 1980; Yoshida, 1994), or by using a buffer that only modifies the external pH and cannot pass the PM (Buffer B: 25 mM MES-BTP pH 5.4). Similar to elicitors and hyphae treatments, Buffer A induced depletion of CSCs from the PM and depolymerization of cortical MTs within 5 min, while Buffer B induced no visible changes in both CesA and MTs density (Supplementary Fig. 3a-d), corroborating that indeed a ∆pH change across the PM caused the observed effects.

### CSC and cortical MT regulation upon biotic stress requires the CC proteins

Our data show a simultaneous response of CSCs and cortical MTs to biotic stress, highlighting their reported interdependence. Considering their physical interaction, we evaluated the role of CSC-MT connecting proteins in response to Fo5176. The first obvious candidates were the CC proteins, as major players in regulating cellulose synthesis and MT dynamics under abiotic stress conditions, such as salt (Endler *et al*, 2015). Therefore, we exposed a *cc1cc2* YFP-CesA6 mChTUA dual-label line (Endler *et al*, 2015), which lacks the two most important CC proteins but is indistinguishable from WT under mock conditions, to Fo5176 hyphae. CSC and cortical MT density at the PM, as well as CSC speed were maintained in *cc1cc2* as compared to WT (Fig. 3 a-d; Supplementary Fig. 4 a and b), indicating that the fast CSCs depletion from the PM and depolymerization of MTs in WT roots upon Fo5176 contact is dependent on the CC proteins. Further supporting these observations, no immediate growth rate reduction was detected for *cc1cc2* when exposed to Fo5176 hyphae (Fig. 3d). To better characterize the role of the CC proteins in plant-microbe interactions at a molecular level, we tested the function of the CC1 MT-interacting domain, recently reported to be essential for plants to regulate MT bundling and dynamics under salt stress (Endler *et al*, 2015). Therefore, we used *cc1cc2* mutant lines complemented with a truncated CC1 version, in which the MT-interacting domain is exchanged to GFP (GFP-CC1ΔN120; (Endler *et al*, 2015)). In this setting, we used GFP-CC1ΔN120 or GFP-CC1 as proxy for CSCs as they co-localize at the PM (Endler *et al*, 2015). No alteration of CSC and MT density as well as no CSC speed reduction could be observed for *cc1cc2* GFP-CC1ΔN120 (Fig. 3e-h; Supplementary Fig. 4c), rendering the plants indistinguishable from mock treated WT plants (Fig. 1b and c; Supplementary Fig. 1a and b). These results demonstrate that the CC protein family does not only participate in plant responses to abiotic but also to biotic stress, but with contrary roles.

**Figure 3:**
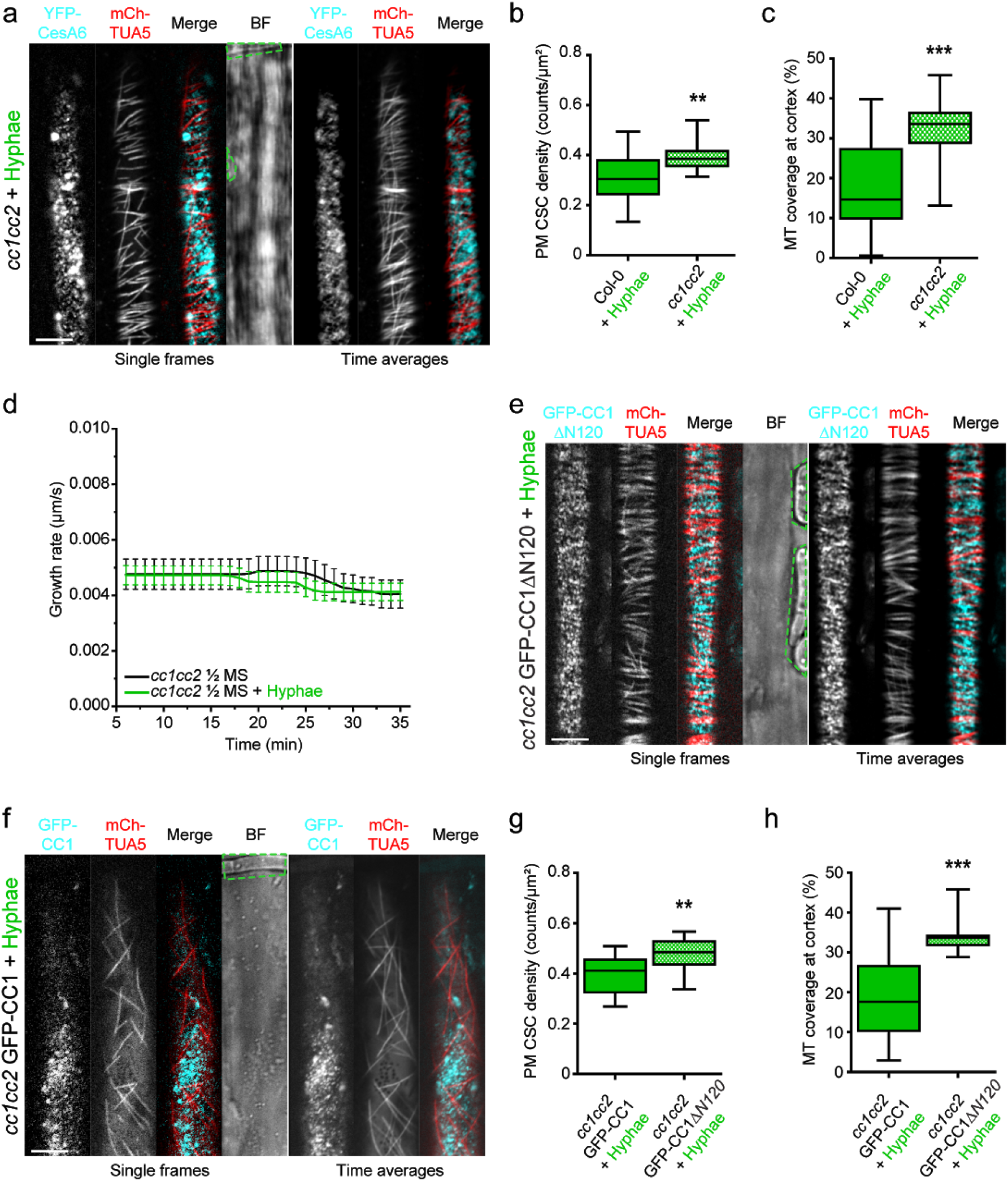
Fo5176 hyphae do not affect the cellulose synthase machinery or the growth rate of *cc1cc2* mutant roots. **a** Representative image of a 5 day-old *cc1cc2* GFP-CesA3 and mCh-TUA5 dual-labeled root epidermal cell upon 5 min of Fo5176 hyphae contact (left panels; single frame, right panels; time average projections). Fo5176 hyphae are highlighted by a green dashed line in the brightfield (BF) channel. Scale bar = 5 μm. **b.** Quantification of GFP-CesA3 density at the plasma membrane of wild-type (Col-0) or *cc1cc2* root cells after Fo5176 hyphae contact as depicted in **a**. Box plots: center lines show the medians; box limits indicate the 25th and 75th percentiles; whiskers extend to the minimum and maximum. N ≥ 20 cells from 15 roots and 3 independent experiments; Welch’s unpaired *t*-test; ** p-value ≤ 0.01. **c** Quantification of microtubule density at the cell cortex of wild-type (Col-0) or *cc1cc2* root cells after Fo5176 hyphae contact as depicted in **a.** Box plots as described in **b.** N ≥ 22 cells from 15 roots and 3 independent experiments; Welch’s unpaired *t*-test; *** p-value ≤ 0.001. **d** Growth rate of *cc1cc2* roots grown in half MS or half MS + Fo5176 hyphae. Average growth rate in half MS: 0.0046 ± 0.0005 μm/s; average growth rate in half MS + Fo5176 hyphae: 0.0045 ± 0.0003 μm/s. Values are mean +/-SEM, N = 17 seedlings from 3 independent experiments. Welch’s unpaired *t*-test; p-value = 0.78. **e** Representative image of a 5 day-old *cc1cc2* GFP-CC1ΔN120 and mCh-TUA5 dual-labeled root epidermal cell upon 5 min of Fo5176 hyphae contact (left panels; single frame, right panels; time average projections). Fo5176 hypha is defined by a green dashed line in the BF channel. Scale bar = 5 μm. **f** Representative image of a 5 day-old *cc1cc2* GFP-CC1 and mCh-TUA5 dual-labeled root epidermal cell upon 5 min of Fo5176 hyphae contact (left panels; single frame, right panels; time average projections). Fo5176 hypha is defined by a green dashed line in the BF channel. Scale bar = 5 μm. **g** Quantification of GFP-CC1 or GFP-CC1ΔN120 (as proxy for CesAs) density at the plasma membrane of *cc1cc2* root cells after Fo5176 hyphae contact as depicted in **e** and **f.** Box plots as described in **b.** N ≥ 14 cells from 8 roots and 3 independent experiments; Welch’s unpaired *t*-test; ** p-value ≤ 0.01. **h** Quantification of microtubule density at the cell cortex of *cc1cc2* GFP-CC1 or *cc1cc2* GFP-CC1ΔN120 root cells after Fo5176 hyphae contact as depicted in **e** and **f.** Box plots as described in **b.** N ≥ 14 cells from 8 roots and 3 independent experiments; Welch’s unpaired *t*-test; *** p-value ≤ 0.001.

### *cc1cc2* is less susceptible to Fo5176 than WT plants

Plant growth rate and the cellulose synthase machinery were obviously affected by Fo5176 contact in a CC and pH-dependent manner. Hence, we questioned if this response was maintained throughout a long term Fo5176 infection. Therefore, we designed an in-plate assay that allows for monitoring both root growth and fungal infection at the same time. To confirm that the fungus was able to reach the xylem in those in-plate conditions and monitor the infection of plant roots, we generated a Fo5176 line harbouring a GFP expression cassette under the control of the Fo5176 SIX1 effector promoter (for *Secreted in Xylem 1*; Fo5176 pSIX1::GFP), similar to what has been done earlier for a tomato-infecting strain (van der Does *et al*, 2008). Indeed, we observed GFP signal in Fo5176 pSIX1::GFP when it reached the xylem of the host vasculature (Fig. 4a) and consequently the sensor allowed us to quantify fungal vasculature penetration events over the progression of the infection. We transferred eight day-old vertically-grown plants to half MS plates covered with Fo5176 pSIX1::GFP spores or mock plates. Three days post fungal inoculation (dpi), WT plants showed a reduction of root growth as compared to mock conditions (Fig 4b and c; Supplementary Fig. 5a and b). *cc1cc2* roots outgrew the WT when treated with Fo5176 pSIX1::GFP in the course of the experiment (Fig. 4b and c), but were indistinguishable from WT under mock conditions (Supplementary Fig. 5a and b). The same was observed for WT Fo5176 treatments (Supplementary Fig. 5c), which confirmed that the Fo5176 virulence is not affected by the pSIX1::GFP construct. In agreement with the observed root growth reduction, WT roots also exhibited the first events of fungal entry into the xylem at three dpi (Fig. 4c), and the vascular penetration rate increased over the course of the experiment. In contrast, the vasculature of *cc1cc2* mutants was barely colonized (Fig. 4c), which is in agreement with the increased growth upon fungal inoculation.

**Figure 4:**
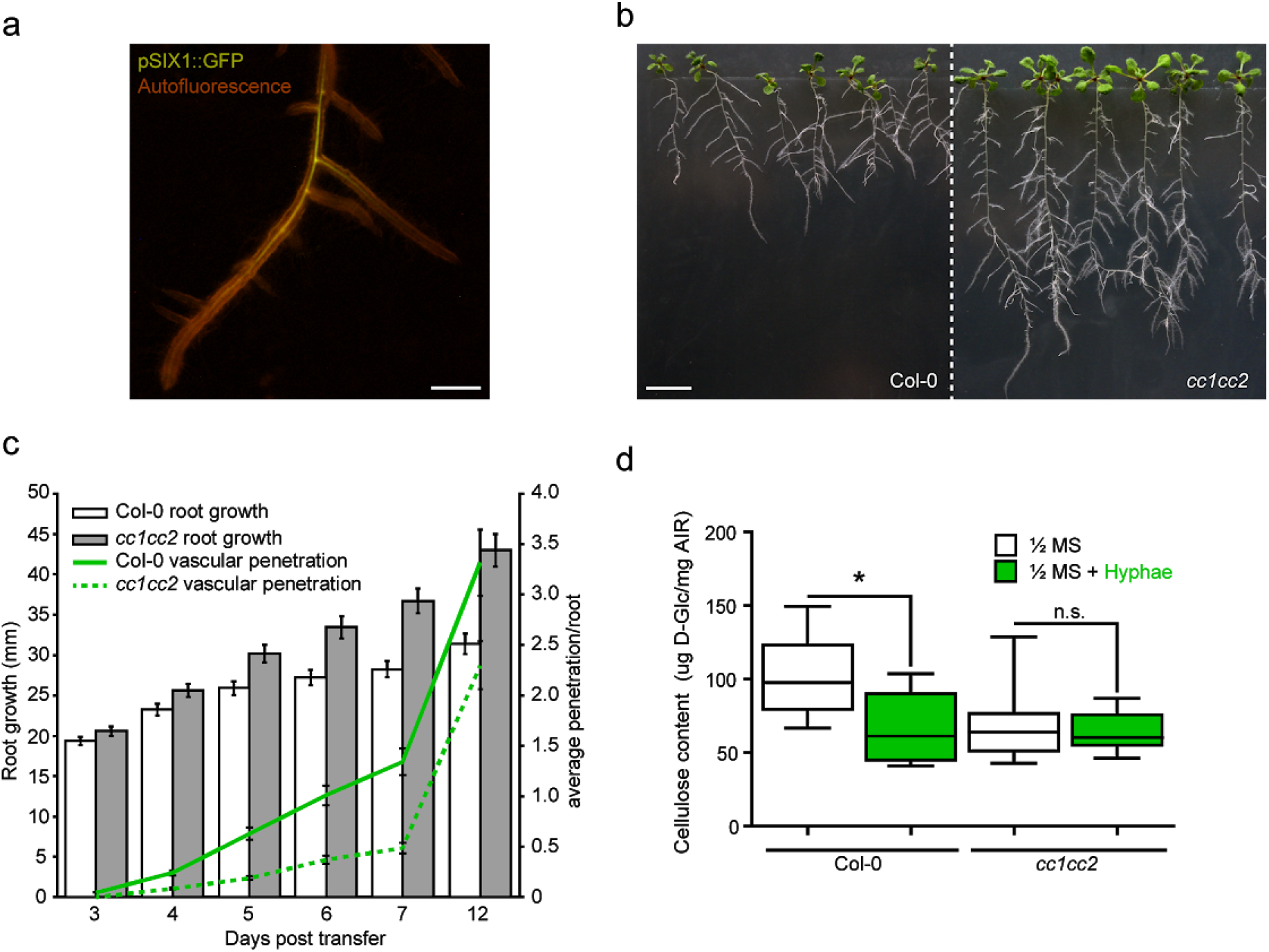
*cc1cc2* mutants are less susceptible than wild-type to Fo5176 root colonization. **a** Arabidopsis root (autofluorescence in red) with a Fo5176 pSIX1::GFP (yellow) colonized xylem. Scale bar = 1 mm. **b** Representative image of wild-type (Col-0) and *cc1cc2* mutant plants 12 days post inoculation with Fo5176 pSIX1::GFP spores. Scale bar = 10 mm. **c** Quantification of root elongation and vascular penetration of wild-type (Col-0; white bars or green line) and *cc1cc2* mutant (grey bars or green dashed line) plants at various days post inoculation with Fo5176 pSIX1::GFP, as depicted in **b**. Values are mean +/-SEM, N ≥ 103 plants from 3 independent experiments. RM two-way ANOVA on root growth: p ≤ 0.001 (genotype), p ≤ 0.001 (time), p ≤ 0.01 (genotype x time). RM two-way ANOVA on vascular penetration rate: p ≤ 0.01 (genotype), p ≤ 0.001 (time), p ≤ 0.001 (genotype x time). **d** Cellulose content of roots grown as in depicted in **b,** represented as μg of crystalline glucose (D-Glc) per mg of dried alcohol-insoluble residue (AIR). Box plots: center lines show the medians; box limits indicate the 25th and 75th percentiles; whiskers extend to the minimum and maximum. N ≥ 5 biological replicates; 2 technical replicates per biological replicate. n.s. = non significant; Welch’s unpaired *t*-test; * p-value ≤ 0.05.

Since the short term effect of hyphae contact on the cellulose synthase machinery could be inhibited by knocking-out the CC proteins and by buffering the media with MES, we also aimed to confirm the influence of the PM ∆pH during long term Fo5176 infection progression. As for the short-term treatment, root growth of Fo5176 pSIX1::GFP treated WT plants was enhanced on buffered media in comparison to non-buffered media (Supplementary Fig. 5d). Simultaneously, the average vasculature penetration rate of Fo5176 pSIX1::GFP was significantly reduced in comparison to penetration rates on non-buffered media (Supplementary Fig. 5d). Finally, we measured the influence of fungal root colonization on the cellulose content of the host. Hence, we adapted a protocol from Yeats & Vellosillo *et al.* to identify all cell wall sugars (Yeats *et al*, 2016a) (Fig. 4d; Supplementary Fig. 5e and f) and were able to reliably distinguish the N-acetylglucosamine of the fungal chitin-based CW from the plant CW derived glucose (Supplementary Fig 5e and f). We observed a significant reduction of cellulose content in WT roots upon Fo5176 pSIX1::GFP colonization, while the fungal infection did not alter the cellulose amount in *cc1cc2* mutants (Fig. 4d).

Based on these results, we conclude that the observed short term fungal effects, i. e. depletion of the cellulose synthase machinery and growth rate reduction, also affect the overall infection process. Furthermore, these effects are repressed in the *cc1cc2* mutant and under buffered media conditions.

### *cc1cc2* exhibits an elevated ΔpH across the PM

The *cc1cc2* mutant did exhibit less sensitivity than WT plants to Fo5176 both in the short and long term (Fig 3 and 4). As we observed a clear influence of plant PM ∆pH changes on the root response to the fungus (Fig 2), we speculated that the pH at the PM interface of *cc1cc2* mutant plants might be altered. Therefore, we introgressed both pH_apo_ and pH_cortical_ sensors into the *cc1cc2* knockout line. *cc1cc2* plants already showed significantly different pH_apo_ and pH_cortical_ when compared to WT plants under mock conditions. We measured an average pH_apo_ over 5 min of 5.21 ± 0.20 (Fig. 5a; WT pH_apo_ 5.42 ± 0.32, Fig. 2h; Values are mean +/-SD, N ≥ 12 seedlings from 3 independent experiments; Welch’s unpaired *t*-test; * p-value ≤ 0.05) and an average pH_cortical_ over 5 min of 7.67 ± 0.38 (Fig. 5b; WT pH_cortical_ 7.07 ± 0.64, Fig. 2i; Values are mean +/-SD, N ≥ 11 seedlings from 3 independent experiments; Welch’s unpaired *t*-test; * p-value ≤ 0.05). This implicates an enhanced proton chemical gradient across the PM in the *cc1cc2* mutant background (WT ΔpH = 1.55, *cc1cc2* ΔpH = 2.45). Unlike WT plants, *cc1cc2* did not show any significant change of pH on both PM sides upon Fo5176 hyphae contact (Fig 5a and b). The altered PM ∆pH of *cc1cc2* roots points towards an elevated PM proton pump activity. Indeed, western blotting corroborated an increased phosphorylation of the penultimate AHA threonine residue in *cc1cc2* mutants as compared to WT (Fig. 5c and d). Furthermore, while the penultimate AHA threonine residue was significantly more phosphorylated upon Fo5176 hyphae contact in WT roots, this was not observed in *cc1cc2* mutant roots (Fig. 5c and d). These results confirm the observed lower pH_apo_ and higher pH_cortical_ measured in mock treated *cc1cc2* root cells (Fig. 5a and b) and the absent adjustment of ∆pH across the PM in response to Fo5176 hyphae (Fig. 5a and b). We measured the same hyperactivation and non-responsiveness to Fo5176 of AHAs in the *cc1cc2* GFP-CC1ΔN120 line (Fig. 5e and f), pointing towards an important role of the CC1 MT- interacting domain for pH regulation at the PM. Hyperactivation of AHAs in both *cc1cc2* and *cc1cc2* GFP-CC1ΔN120 mutants was additionally confirmed by measuring the acidification process of an alkaline growth media, which both lines acidified significantly faster than WT plants (Supplementary Fig. 6a). Confirming the higher PM ΔpH of *cc1cc2*, both CC-impaired lines were more sensitive to low amounts of hygromycin B as compared to WT (Supplementary Fig. 6b-e), which is primarily internalized through the energy generated by the proton chemical gradient (Haruta *et al*, 2010). To assess if hyperactive AHAs in *cc1cc2* also play a role in the long term Fo5176 infection process, we measured the media pH of liquid plant cultures treated with Fo5176 pSIX1::GFP spores at various dpi. We observed an alkalinization of the media in the Fo5176 infection process (Fig. 5g and h), confirming previous data reported in the pathosystem tomato-*Fusarium oxysporum f. sp. lycopersici* (Masachis *et al*, 2016). But, the process was significantly delayed for media containing *cc1cc2* plants (Fig. 5g and h), implicating that the hyperactivation of AHAs leads to a delay in alkalinization and less susceptibility to Fo5176.

**Figure 5:**
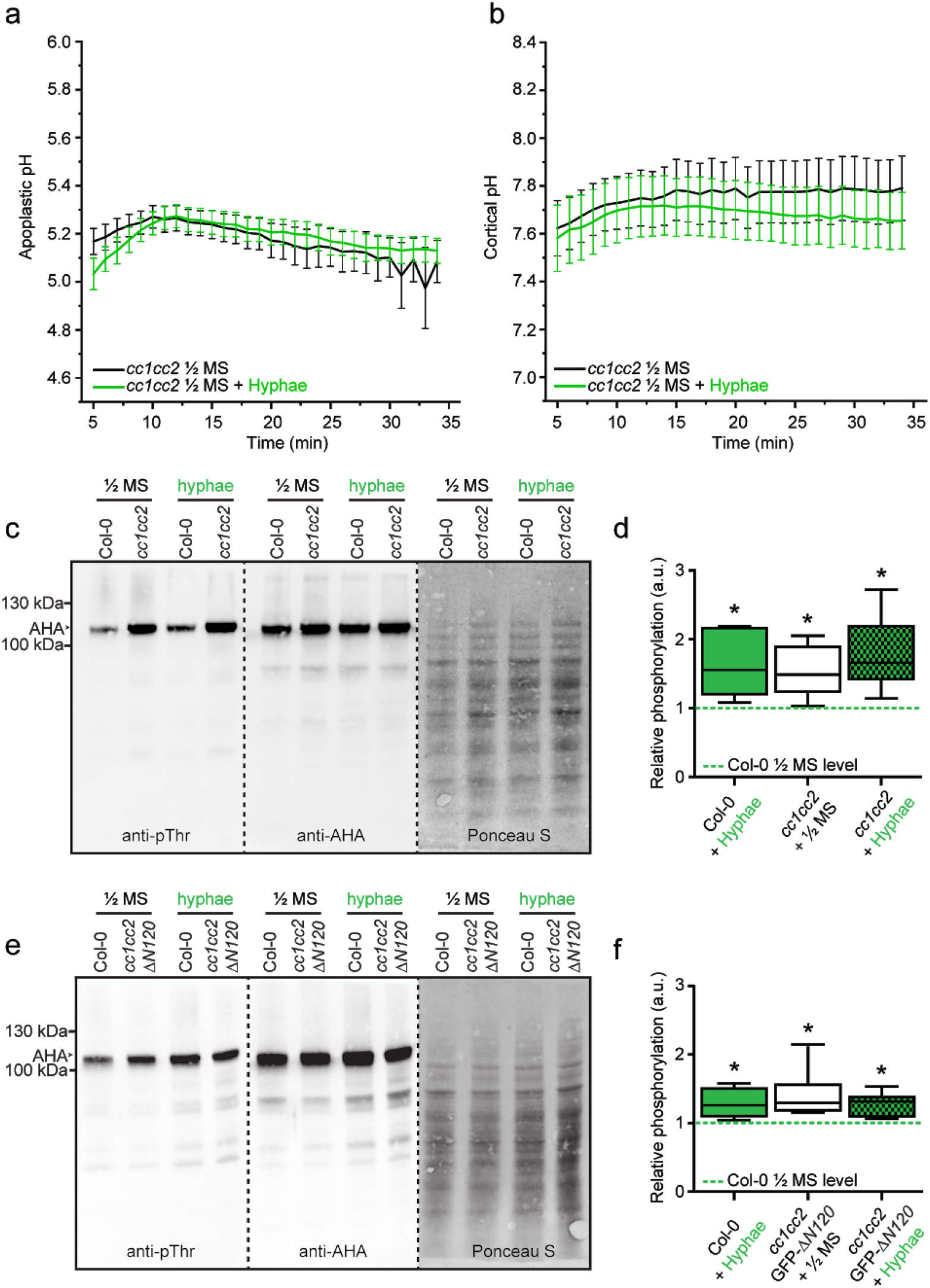
*cc1cc2* mutants exhibits an acidic apoplast, alkaline cortex and high AHA phosphorylation state, which are not affected by Fo5176 contact. **a** Apoplastic pH variation of *cc1cc2* roots expressing the pH_apo_ sensor SYP122-pHusion over time, either in half MS or half MS + Fo5176 hyphae. Values are mean +/-SEM, N = 17 seedlings from 3 independent experiments. Welch’s unpaired *t*-test; p-value = 0.84. **b** Cortical pH variation of *cc1cc2* roots expressing the pH_cortical_ sensor pHGFP-Lti6b over time, either in half MS or half MS + Fo5176 hyphae. Values are mean +/-SEM, N ≥ 12 seedlings from 3 independent experiments. Welch’s unpaired *t*-test; p-value = 0.67. **c** Western blots showing chemiluminescent signals of anti-pThr or anti-AHA incubated membranes loaded with Arabidopsis wild-type (Col-0) or *cc1cc2* root samples. Roots were treated for 8 min with either half MS or half MS + Fo5176 pSIX1::GFP hyphae. The Ponceau S panel shows total protein content. The AHA band used for quantification is highlighted with an arrowhead. Dashed line separates different treatments of the same membrane. **d** Quantification of AHA phosphorylation status from western blots as shown in **c**, calculated by obtaining the signal intensity ratio of anti-pThr in respect to anti-AHA and normalized against the Col-0 half MS level (green dashed line). Box plots: center lines show the medians; box limits indicate the 25th and 75th percentiles; whiskers extend to the minimum and maximum. N = 5 independent experiments. Welch’s unpaired *t*-test in comparison to Col-0 mock conditions; * p-value ≤ 0.05. **e** Western blots showing chemiluminescent signals of anti-pThr or anti-AHA incubated membranes loaded with *Arabidopsis* wild-type (Col-0) or *cc1cc2* GFP-CC1CC1ΔN120 root samples. Roots were treated for 8 min with either half MS or half MS + Fo5176 pSIX1::GFP hyphae. The Ponceau S panel shows total protein content. The AHA band used for quantification is highlighted with an arrowhead. Dashed line separates different treatments of the same membrane. **f** Quantification of AHA phosphorylation status from western blots as shown in **e**, calculated by obtaining the signal intensity ratio of anti-pThr in respect to anti-AHA and normalized against the Col-0 half MS level (green dash line). Box plots as described in **d**. N=5 independent experiments. Welch’s unpaired *t*-test in comparison to Col-0 mock conditions; * p-value ≤ 0.05.

## Discussion

Under natural conditions, plants are constantly exposed to abiotic and biotic stresses. Studying how plants cope with these factors is therefore of utmost importance in times of uncertain environmental and climate changes. Here, we outline a detailed analysis of early and long term plant cellular responses to pathogen-induced stress at the host PM interface.

Our work demonstrates that the plant cellulose synthesis machinery, composed of PM-active CSCs and cortical MTs, is remarkably sensitive to biotic stress. The cortical MT array has been reported to concentrate on the contact site of plant cells with hyphae or a microneedle (Hardham *et al*, 2008), while a low degree of de-polymerization was observed 75 min after treatment with VD toxin from the fungus *Verthicillium dahliae* (Yao *et al*, 2011). We show an unexpected, rapid depolymerization of MTs within 5 min upon contact with Fo5176 hyphae and Fo5176-derived molecules (Fig 1). Moreover, our assays showed a simultaneous decrease of density and speed of PM-located CSCs (Fig 1) that likely not stems from MT depolymerization, as the CSC status is not affected when MTs are chemically depolymerized (Gutierrez *et al*, 2009; Endler *et al*, 2015). We found that the impairment of the cellulose synthesis machinery was coordinated with a reduction of cell elongation (Fig. 1) and a modulation of AHA phosphorylation state (Fig 2). Negative regulation of AHAs by phosphorylation has been reported in Arabidopsis cell cultures exposed to the bacterial elicitor flg22 (Nühse *et al*, 2007; Benschop *et al*, 2007). In contrast, our data show a fast AHA activation in intact Arabidopsis roots (Fig 2j) that we could directly link to real changes of pH at both sides of the PM with newly developed pH sensors (Fig 2). Our work adds a detailed layer of spatial resolution to the analysis of subcellular pH and demonstrates the high level of cell compartmentalization, as ΔpH changes were restricted to the PM interface and did not affect the pH_cyto_, probably due to a buffering effect of the tonoplast (Pittman, 2012). The pH_apo_ and pH_cortical_ sensors are novel molecular tools, which allow for reliable measurement of apoplastic and PM-cortical pH with little intracellular background (Gjetting *et al*, 2012; Martinière *et al*, 2018). They enabled us to perform real time measurements of fast, fungal-induced acidification of the root cell apoplast and ultimately cortex, which derived from the upregulation of AHAs (Fig 2). This novel insight into pH regulation in response to stress helps to explain previous observations of fast media acidification close to microbe exposed roots (Felle *et al*, 2009). Our data indicate that the fast PM ΔpH variation in response to Fo5176 triggered the observed CSCs and cortical MT alterations, as well as a reduction of cell elongation, since all these cellular reactions could be blocked with 5 mM MES buffer (Fig. 2 and Supplementary Fig. 2). In addition, the CSC-MT disturbance by Fo5176 could be mimicked by exposing roots to a buffer that changes the PM ΔpH without any abiotic or biotic stress (Supplementary Fig. 3). The influence of intracellular pH changes on the cortical MT array has recently been reported in Chlamydomonas (Liu *et al*, 2017). These and our observations point towards a conserved mechanism in algae and land plants in which stress-induced cellular pH changes alter the MT network. Various complementary scenarios could explain the fast response of MTs to pH changes driven by stress. PM-linked MTs might sense changes in PM lipid composition, which is influenced by the PM ΔpH (Platre *et al*, 2018) or MTs might be linked to certain ion-channels whose activity is potentially regulated by pH changes at the PM interface and causes activation of downstream signaling cascades (Prager-Khoutorsky *et al*, 2014). In agreement with our observed short term effects, Fo5176 root infection led to a reduction of cellulose content (Fig 5), confirming the expected consequence of pathogenic root colonization.

The CC proteins and more specifically the CC1 MT-binding domain are indispensable for the rearrangement of the CSC-MT machinery in response to salt stress (Endler *et al*, 2015; Kesten *et al*, 2018). Consequently, mutants affected in CC function show stunted growth on salt containing media (Kesten *et al*, 2018). Here, we show that the same CC1 domain is directly involved in the regulation of the CSC-MT machinery upon biotic stress. Contrary to the phenotype on salt, *cc1cc2* was less susceptible than control plants to Fo5176 (Fig 3 and 4). Interestingly, salt stress was reported to alkalinize the apoplast and the cytosol (Gao *et al*, 2004; Geilfus, 2017), which reflects the exact opposite response of what we observed upon fungal contact: a fast apoplastic and cortical acidification (Fig. 2). This highlights the inverse effect of biotic and abiotic stress on plant cellular pH modulation at the PM and might explain the opposite phenotypes of *cc1cc2* on abiotic and biotic stress. The CC1 MT-binding domain is involved in PM pH regulation by controlling AHA phosphorylation state, as the proton pumps showed a permanently elevated basal activity in *cc1cc2* and *cc1cc2* GFP-CC1ΔN120, which did not get further activated in response to the fungus (Fig. 4). Consequently, *cc1cc2* exhibited a more acidic apoplast and basic cortical side of the PM (Fig. 3) and was able to acidify a basic growth media faster than WT plants (Fig. S5). Similar to the short-term response, root growth inhibition and reduction of cellulose deposition caused by the fungal infection were CC-dependent (Fig. 4). At the same time, the ability of Fo5176 to colonize the xylem was reduced *cc1cc2* (Fig. 4) and in wild-type plants grown on buffered media (Supplementary Fig. 5). Thus, a high PM ΔpH might be positive for plant defense, but negative for adaptation to salt stress. Host tissue alkalinization has been shown to be needed for *F. oxysporum* infection of tomato roots (Masachis *et al*, 2016; Stegmann *et al*, 2017; Sánchez-Rangel *et al*, 2018). In addition, the activation of PM H^+^-ATPases seems to potentiate plant defense mediated by resistance genes, while apoplastic alkalinization induced by microbe-associated molecular patterns has been reported to precede cell death (Elmore & Coaker, 2011). The role of pH in plant-microbe interaction is a complex question, as lasting apoplastic alkalinization, on the other hand, has been suggested to support plant defense against pathogens, e.g. via plant CW stiffening (Felle *et al*, 2005). Environmental pH has furthermore been reported to modulate both plant and microbe gene expression (Masachis *et al*, 2016; Stegmann *et al*, 2017; Sánchez-Rangel *et al*, 2018) and to influence the activity of microbe-derived plant CW degrading enzymes (Roncero *et al*, 2000; Reverchon & Nasser, 2013). Our data strongly suggest that changes in PM ΔpH caused by both salt (Gao *et al*, 2004; Geilfus, 2017) and Fo5176 (Fig 3-5) are CC-dependent and a main stress signal that generates cortical MT depolymerization and CSC depletion from the PM. In consequence, this might lead to constraint of cellulose synthesis in both cases. Our work offers new molecular tools to understand the precise role of spatiotemporal changes in the cellular pH of plants in response to stress, its coordination with cellulose synthesis and cell expansion and how the CC proteins are coupled to this.

In conclusion, our data extends the current view of plant adaptation to the environment and indicates that the PM proton chemical gradient is modulated in a CC-dependent manner. The gradient, therefore, might act as a decisive starting point for both biotic and abiotic signalling cascades. Furthermore, we show a new set of proteins that connects growth and defense, with opposite functions upon biotic and abiotic stress, which should be kept in mind in the attempt to create stress tolerant plants.

## Materials & Methods

### Plant material and growth

*Arabidopsis thaliana* (Col-0) lines expressing pUB10::SYP122-pHusion and pUB10::pHGFP-Lti6b were transformed according to standard procedures (Hellens *et al*, 2000). Transgenic lines were isolated on plates containing 0.5 x MS (Duchefa), 0.5 % sucrose, pH 5.8 (KOH) and 0.55 % phytoagar (Duchefa) supplemented with either 11.25 µg/ml sulfadiazine for pUB10::SYP122-pHusion or 50 µg/ml hygromycin B for pUB10::pHGFP-Lti6b. Transgenic lines were screened for 3:1 segregation of the resistance marker and fluorescence intensities of the respective sensors. The pH_cyto_ sensor line, pUB10::pHGFP and the *cc1cc2*, *cc1cc2* GFP-CC1 and *cc1cc2* GFP-CC1ΔN120 lines were published previously (Endler *et al*, 2015; Fendrych *et al*, 2014).

### Fungal strains, culture conditions and elicitor mix preparation

*Fusarium oxysporum* Fo5176 was used throughout this study. Strain culture and storage was performed as described earlier (Di Pietro *et al*, 2001). Fungal elicitor mix was prepared based on a method published previously (Baldrich *et al*, 2014) with the following modifications: Fo5176 was grown in liquid Potato Dextrose Broth (PDB) at 27°C in the dark for 5 days. The culture was filtered through miracloth and washed thoroughly with water to remove excess PDB. Mycelia were harvested, frozen in liquid nitrogen and lyophilized until dry. Dried mycelia were ground to powder in a Geno/Grinder (SPEX SamplePrep, U.S.A). The powder was diluted with distilled water to obtain a concentration of 30 mg/ml. The slurry was autoclaved for 15 min at 121°C, aliquoted and stored at −20°C.

### Generation of Fo5176 hyphae for short treatments of roots

1 ml half MS + 1% sucrose containing 10^7^ Fo5176 or Fo5176 pSIX1::GFP spores was shaken horizontally overnight at 150 rpm in a 2 ml tube on a rotating device. Germinated spores were spun down at 2,000 g for 5 min. Supernatant was discarded and hyphae were washed 3 times with half MS (pH 5.7) to remove excess sucrose.

### Constructs

pUB10::SYP122-pHusion and pUB10::pHGFP-Lti6b constructs were generated and assembled using GreenGate (GG) cloning(Lampropoulos *et al*, 2013).

For pUB10::SYP122-pHusion, the full-length coding sequence (CDS) of SYP122 (Syntaxin of Plants 122; At3g52400) was amplified from *Arabidopsis thaliana* Col-0 cDNA with primers 1 and 2 listed in Supplementary Table 1, attaching Eco31I recognition sites and specific GG-overhangs. The Stop codon was removed from the SYP122 CDS and the resulting 1056-bp fragment was subcloned into pGGC000 (Lampropoulos *et al*, 2013). The CDS of pHusion was amplified from p16-SYP61-pHusion (Luo *et al*, 2015) with primers 3 and 4 listed in Supplementary Table 1 and the resulting 1439-bp fragment was subcloned into pGGD000 (Lampropoulos *et al*, 2013). The final construct was assembled in a GG reaction from modules listed below.

- pGGA006 (UBIQUITIN10 promoter; (Lampropoulos *et al*, 2013))
- pGGB003 (B-dummy; (Lampropoulos *et al*, 2013))
- pGGC-SYP122 (SYP122 CDS)
- pGGD-pHusion (pHusion CDS)
- pGGE001 (RBCS terminator from pea; (Lampropoulos *et al*, 2013))
- pGGF012 (pMAS:SulfRm:t35S; (Lampropoulos *et al*, 2013))
- pGGZ001 (Vector backbone with plant resistance at RB; (Lampropoulos *et al*, 2013)

To generate pUB10::pHGFP-Lti6b, the CDS of pHGFP was amplified from proUBQ10:pHGFP (Fendrych *et al*, 2014) in two parts, to remove an internal Eco31I site by introducing a silent point mutation. Eco31I recognition sites and specific GG-overhangs were attached to amplicons with primers 5-8 listed in Supplementary Table 1. The internal Eco31I site and the Stop codon was removed from pHGFP and the resulting 676-bp and 99-bp fragments were subcloned into pGGC000 (Lampropoulos *et al*, 2013). The CDS of Lti6b was amplified from PM-YC3.6-Lti6b (Krebs *et al*, 2012) with primers 9 and 10 listed in Supplementary Table 1 and the resulting 195-bp fragment was subcloned into pGGD000 (Lampropoulos *et al*, 2013). The final construct was assembled in a GG reaction from modules listed below.

- pGGA006 (UBIQUITIN10 promoter; (Lampropoulos *et al*, 2013))
- pGGB003 (B-dummy; (Lampropoulos *et al*, 2013))
- pGGC-pHGFP (pHGFP CDS)
- pGGD-Lti6b (Lti6b CDS)
- pGGE001 (RBCS terminator from pea; (Lampropoulos *et al*, 2013))
- pGGF005 (pUBQ10:HygrR:tOCS; (Lampropoulos *et al*, 2013))
- pGGZ003 (Vector backbone with plant resistance at LB; (Lampropoulos *et al*, 2013)

For plant transformation pUB10::SYP122-pHusion and pUB10::pHGFP-Lti6b were transformed into *Agrobacterium tumefacien*s ASE strain harbouring the pSOUP plasmid.

To obtain the Fo5176 pSIX1::GFP line, the pRW2h binary vector (Houterman *et al*, 2008) was digested (XbaI/HindIII) and a three point ligation was performed using two inserts, which were amplified with primers 11-18 listed in Supplementary Table 1: (1) the 1kb region downstream of the Fo5176-SIX1 ORF, digested with XbaI/EcoRI, and (2) the Herpes Simplex Virus thymidine kinase (HSVtk) gene under the control of the *Cochliobolus heterostrophus* glyceraldehyde-3-phosphate dehydrogenase (ChGPD) gene promoter and the *Neurospora crassa* β-tubulin gene terminator (EcoRI/HindIII). Since this vector was meant for use in a knock-out experiment of SIX1, HSVtk was inserted as a conditional negative selection marker against ectopic transformants (Khang *et al*, 2005). Subsequently, the 1kb region upstream of Fo5176-SIX1 was inserted into the vector using PacI/ClaI, and the GFP coding sequence was inserted next to this promoter region using primers with ClaI and KpnI linkers. The plasmid was inserted into *F. oxysporum* Fo5176 by *Agrobacterium* mediated transformation as described previously (Takken *et al*, 2004). One ectopic transformant was selected for having low *in vitro* and higher *in planta* GFP expression.

### Spinning disc live cell-imaging and data processing

5 day old *A. thaliana* seedlings grown upright on non-buffered, solid half MS media (pH 5.7) in a 16h light/8h dark cycle were used for all experiments. Plant roots were covered with a 1% agarose cushion as described previously (Gutierrez *et al*, 2009). Treatments were added to coverslips before plant transfer and roots were subsequently placed directly into the treatment solution. 10 µl half MS (pH 5.7) containing approximately 10^4^ Fo5176 hyphae (originated from overnight germination of Fo5176 spores, see above) or an elicitor mix in a 1:3.66 dilution with half MS were used as treatment. The whole process (transferring to the microscope and adjustments) took approximately 5 min.

XFP-tagged proteins were imaged with a CSU-W1 Yokogawa spinning disc head fitted to a Nikon Eclipse Ti-E inverted microscope with a CFI PlanApo × 100 N.A. 1.40 oil immersion objective, an EM-CCD ImageEM 1K (C9100-14) (Hamamatsu Photonics, Japan), and a ×1.2 lens between the spinning disc and camera. GFP was imaged using a 488 nm solid state diode laser and a 525/50 nm emission filter, RFP was detected with a 561 nm solid state diode laser and a 609/54 nm emission filter. Alternatively, a CSU-W1 Yokogawa spinning disc head fitted to a Nikon Eclipse Ti-E inverted microscope with a CFI PlanApo × 100 N.A. 1.40 oil immersion objective, two iXon Ultra EM-CCD cameras (Andor, GB), and a ×1.2 lens between the spinning disc and camera was used. For this system, GFP was imaged using a 488 nm solid state diode laser and a 525/50 nm emission filter, RFP was detected with a 561 nm solid state diode laser and a 630/75 nm emission filter and YFP was detected with a 515 nm solid state diode laser and a 535/30 nm emission filter. Time lapse images were processed and analyzed with Fiji (Schindelin *et al*, 2012). Drifts were corrected by using the plugin StackReg or MultiStackReg in cases where two channels were imaged (Thevenaz *et al*, 1998). Backgrounds were subtracted by the “Subtract Background” tool (rolling ball radius, 30–50 pixels). To quantify CesA velocities three frames were averaged by “WalkingAverage” and kymograph analysis was performed with the kymograph tool of FIESTA (Ruhnow *et al*, 2011).

### CesA and microtubule density measurements

Both methods were described earlier (Endler *et al*, 2015). MT density measurements were done following the basic principle of the method described above but it was transferred to Fiji. Cell boundaries were detected by applying a Gaussian kernel (sigma =1.33 μm) into each image and applying a various thresholds with the Otsu algorithm (Otsu, 1979). Instead of using the mentioned Sobel edge-detection algorithm, a Laplacian image was generated (Gaussian kernel with sigma = 1.5 μm) using FeatureJ (Erik Meijering, Biomedical Imaging Group, EPFL Lausanne). Various thresholds, based on signal to noise ratio of each individual image series, were applied to detect most microtubules and least noise pixel. The total area of microtubules was then set in relation to the cell area, which resulted in microtubule density.

### Live cell ratiometric pH sensor imaging including flat and dark field correction and data processing

5 day-old *A. thaliana* seedlings grown upright on non-buffered, solid half MS media (pH 5.7) in a 16h light/8h dark cycle were transferred to imaging chambers as described earlier (Krebs & Schumacher, 2013). 400 µl of half MS (pH 5.7) or half MS containing 5 mM MES (pH 5.7) were used as imaging media. 150 µl fungal elicitor mix were added at the indicated time points through the opening in the lid of the imaging chambers. For hyphae treatment and growth measurements, as a minor modification, the seedlings were not glued to the surface of a coverslip but placed on top of a 1% agarose cushion as described previously (Gutierrez *et al*, 2009). 10 µl of half MS containing approximately 10^5^ Fo5176 hyphae (originated from overnight germination of Fo5176 spores, see above) were spread across the agarose sandwich and air dried for 5 min before seedlings were placed on top. Imaging was started immediately afterwards and the whole process (transferring the chamber to the microscope and adjustments) took approximately 5 min.

Imaging of plant roots was performed on a Leica TCS SP8-AOBS (Leica Microsystems, Germany) confocal laser scanning microscope equipped with a Leica 10x 0.3NA HC PL Fluotar Ph1 objective. pHusion was excited simultaneously, where GFP was excited with 488 nm and detected between 500-545 nm and mRFP was excited with 561 nm and detected between 600 to 640 nm. pHGFP-Lti6b and free pHGFP were excited sequentially, where GFP was first excited with 405 nm and detected between 500-545 nm and then excited with 488 nm and detected between 500-545 nm. HyD detectors were used for all image acquisitions. Image settings were kept identical for calibration and pH measurements with offsets being deactivated. Except for the standards, all images were collected as an XYt series for 30 min with a time frame of 1 min.

To collect standard curves for the pH sensors, 6-12 seedlings each were incubated for 15 min in a buffer series with pH values between 4.8-8.0. The buffers contained 50 mM 4-(2-hydroxyethyl)-1-piperazineethanesulfonic acid (HEPES) (pH 6.8-8.0) or 50 mM 2-(N-morpholino)ethanesulfonic acid (MES) (pH 5.2-6.4) and 50 mM ammonium acetate. The pH was adjusted to the desired value by adding Bis-tris propane (BTP). If necessary, the pH 5.2 buffer was additionally adjusted by adding a small amount of diluted HCl. Buffer pH 4.8 was composed of 22 mM citric acid, 27 mM trisodium citrate and 50 mM ammonium acetate. pH was adjusted with small amounts of KOH and HCl. After incubation, seedlings were placed between a microscope slide and coverslip and imaged as described above.

Flat field images were collected as described previously (Model & Burkhardt, 2001; Model, 2006). The procedure was done for each detector-objective combination. Dark images were acquired by setting the lightpath of the microscope to the eyepiece and therefore blocking all light to reach the detector. Subsequently, a time series of 50 images was acquired and averaged using the Z-projection tool of Fiji in average intensity mode. Flat field images were acquired by using the following dyes and concentrations to prepare imaging slides:

- 7-Diethylamino-4-methylcoumarin (Sigma D87759-5G, 50 mg/ml in DMSO)
- Fluorescein sodium salt (Sigma 46960-25G-F, 100 mg/ml in H_2_O)
- Rose bengal (Sigma 198250-5G, 100 mg/ml in H_2_O)
- Brilliant Blue FCD (Sigma 80717-100MG, 100 mg/ml in H_2_O)

A drop of the specific dye solution was placed between slide and coverslip. A uniform dye area was selected and the focus was set to the area just below the coverslip. Subsequently, 30 different spots were imaged with settings that used the whole range of the detector (i.e. no over- or underexposure). These images were averaged using the Z-projection tool of Fiji in median intensity mode. The image was converted back to 8-bit with disabled scaling. All acquired XYt image series and standards were finally automatically corrected with a self written Fiji macro, which did the following modifications:

1. dark images were subtracted from each single image of a series by using the Fiji plugin Calculator Plus.
2. each single image of the resulting dark corrected series was divided by the median flat field image with Calculator Plus.
3. each single image of the resulting dark and flat field corrected image series was multiplied with the average intensity of the corresponding flat field image using Calculator Plus to raise intensity back to appropriate values

Subsequently, ratio calculations (488 nm/561 nm for pHusion, 405nm/488 nm for pHGFP) were performed on the corrected image series. The calculations were performed automatically with a self written Fiji macro, which did the following operations:

1. a maximum projection over the whole image series was generated to estimate an average root position over time and correct for root growth and (slight) drifting of the root
2. root boundaries were detected on this maximum projection by applying a Gaussian kernel (sigma =1.33 μm) into each image and applying various thresholds with the Otsu algorithm (Otsu, 1979). All following operations were performed in these boundaries
3. for each channel, a specific LUT was applied with the set Min and Max function, which was kept constant for all corresponding image series in this manuscript. At a minimum, the range was set to 1-254 to remove all black (0) and oversaturated (255) pixel
4. the macro then iterated through the slices and channels of the image series and automatically divided the mean gray value of the first channel by the corresponding second channel (e.g. 488 nm/561 nm) in each slice and exported the ratio over time

Standard curves were calculated using sigmoidal regression (4PL fitting algorithm) with Prism (version 8, Graphpad, USA). The pH over time of the image series was subsequently calculated based on the equations of the standard curve. The replicates of individual time series were averaged and visualized with the multiple curve averaging function of OriginPro 2019 (OriginLab), USA).

### Root growth analysis

5 day-old seedlings were mounted into an imaging chamber and treated as described above in the “live cell ratiometric pH sensor imaging” section and experiments were undertaken in the same manner. Root growth was measured by performing a time-phased image subtraction with a phase shift of one frame. The detailed method was described earlier (Lindeboom *et al*, 2013; Endler *et al*, 2015). The resulting “root growth fronts” were transformed into kymographs by drawing a segmented line (line width = 2 pixel) through the tip of the root over time, followed by the kymograph plugin of Fiji. Positions of the root tip over time were extracted from the kymographs by following the kymographs with a segmented line and finally exporting the coordinates with the “save as XY Coordinates” function of Fiji.

### Plate infection assay

1×10 cm Whatman paper strips were heat sterilized, and two strips placed on each 12×12 cm square plate containing 50 ml solid half MS media. 6-10 sterilized and stratified *A. thaliana* seeds were placed on each strip. The plants were grown vertically for 8 days in a 16h light/8h dark cycle. 200 µl of water containing 2×10^6^ Fo5176 or Fo5176 pSIX1::GFP spores were equally spread on a 12×12 cm square plate containing 50 ml solid half MS media. One Whatman paper strip with 8 day-old *A. thaliana* plants was transferred to the plate. The plate was scanned each day to assess root growth. Root length was measured with Fiji. Fo5176 pSIX1::GFP penetration of the root vasculature was assessed with a Leica M205 FCA fluorescent stereo microscope, equipped with a long pass GFP filter (ET GFP LP; Excitation nm: ET480/40x; Emission nm: ET510 LP). Vascular penetration/infection was counted when clear GFP signal was observed.

### Plant CW analysis

10 day-old light-grown *Arabidopsis* roots, under either mock or *F. oxysporum*-infected conditions, were harvested, flash-frozen, and subsequently lyophilized using a freeze-dryer (Kühner, Alpha 2-4). Lyophilized roots were ground with glass beads (2.85-3.45 mm beads, Roth, Article number A557.1) using a tissue homogenizer (Retsch MM301). Starch degradation was performed as previously described (Hostettler *et al*, 2011). Alcohol-insoluble residue (AIR) production, subsequent sample preparation, hydrolysis, and data analysis for monosaccharide quantification was performed as previously described (Yeats *et al*, 2016b, 2016a) with some modifications. Prior to hydrolysis, 150μg of sedoheptulose (CarboSynth, Item Number MS139006) was added to each sample as an internal standard. 10μl of 1:10 dilutions of hydrolyzed samples were measured on a Dionex ICS-5000 using a ThermoFischer Scientific CarboPac PA20 column (3 x 150 mm, Product Number 060142) and accompanying CarboPac PA20 guard column (3×30 mm, Product Number 060144). Eluents for HPLC analysis were prepared as follows: Eluent A: water, Eluent B: 50 mM NaOH, Eluent C: 100 mM Na-Acetate, 100 mM NaOH, Eluent D: 200 mM NaOH. All eluents were purged with and maintained under helium gas as described by the manufacturer. Column temperature was maintained at 36°C as well as a constant eluent flow of 0.4 ml/min of the following gradient to elute monosaccharides: 0-18 min 4.8 % B, 95.2 % A; 18-20 min linear increase to next condition; 20-30 min 50 % D, 50 % A; 30-40 min linear increase to next condition; 40-56 min 100 % C; 56-56.1 min linear decrease to 50 % D; 56.1-60 min 50 % D; 60-60.1 min linear decrease to next condition; 60.1-80 minutes 4.8 % B, 95.2 % A to equilibrate column back to starting conditions. All standard curve and hydrolysis sample peaks were integrated using Chromeleon 8.0 software.

### Quantification of AHA phosphorylation levels

Roots from 10 day-old plants were transferred to solid half MS media plates and treated for 8 minutes with 50 µl half MS or with 50 µl of half MS containing approximately 5×10^5^ Fo5176 or Fo5176 pSIX1::GFP hyphae (originated from overnight germination of Fo5176 spores, see above). Roots were collected and grinded with a pestle in 30 µl pre-heated (65 °C) Laemmli sample buffer (Laemmli, 1970) and solubilized for 4 min. The homogenates were centrifuged at room temperature (10,000 x *g* for 5 min) and 20 µl of the supernatant were loaded onto 4-12% (w/v) acrylamide gradient gels (Expedeon, GB). SDS-PAGE and protein transfer to nitrocellulose membranes were performed with a Trans-Blot Turbo Transfer System (BioRad, USA) according to the manufacturer’s protocol. The amount of transferred protein was analyzed by staining the membrane with Ponceau S Staining Solution (0.1% (w/v) Ponceau S in 5% (v/v) acetic acid) 10 min and washing with distilled water to remove the background. Two polyclonal AHA antibodies were subsequently used: first, one against the region including the phosphorylated penultimate threonine residue (anti-pThr), then one against the conserved catalytic domain of the AHA (anti-AHA) (Hayashi *et al*, 2010). A goat anti-rabbit IgG conjugated to horseradish peroxidase (AC2114, Azure biosystems, U.S.A) was utilized as a secondary antibody and the chemiluminescence from the horseradish peroxidase reaction with a chemiluminescence substrate (WesternBright ECL, Advansta, U.S.A) was detected using the Chemidoc Touch Image System (BioRad, USA). The membrane was stripped with 0.5M Glycine (pH 2.2) and reused with the anti-AHA antibody. The signal intensity of bands was analyzed using the “Gel function” of Fiji and processed as described previously (Kesten *et al*, 2016). The signal intensity of the anti-phThr antibody was normalized with respect to the intensity of the anti-AHA signal. Data was finally normalized to signal intensity ratios of Col-0 in mock conditions.

### Measurement of media pH

Plants were grown on half MS (pH 5.7) plates for ten days in a 16h light/8h dark cycle prior to transfer to liquid half MS media with different pH and treatments. Media pH was monitored with a pH Microsensor from Mettler Toledo Inlab (U.S.A). The proton secretion assay was performed by transferring one plant into 1 ml half MS (pH 6.45) and the change of media pH was measured over 6 hours. The pH changes in the media in response to Fo5176 were assessed by transferring one plant into 1 ml liquid half MS (pH 5.7), inoculated with 50 µl half MS containing approximately 5×10^5^ Fo5176 hyphae (originated from overnight germination of Fo5176 spores, see above) and the pH of the media was monitored until the death of the plants (6 dpi).

### Hygromycin B sensitivity assay

Plants were grown on half MS plates for seven days prior to transfer to plates containing half MS supplemented with or without 5 µg/ml of Hygromycin (Roth, Germany). After seven days, the plates were scanned and root growth was quantified by employing the Fiji software.

### Statistical analysis and experimental design

For statistical analyses, Welch’s unpaired *t*-test, repeated measures two-way ANOVA, or mixed effects model analyses were performed using GraphPad Prism 8. A p-value of <0.05 was considered as statistically significant. Statistical methods and the resulting p-values are defined in the corresponding figure legends. If the fluorescent ratios of the pH sensors (pHusion, pHGFP) were measured to be outside of the standard curve range, they were excluded from the analysis. If samples drifted during image acquisition and this could not be corrected as described above, they were excluded from the analysis.

## Acknowledgments

We are grateful to G. Sancho-Andrés, B. Pfister, and P. van Dam for technical support. We thank S. Persson for the *cc1cc2*, *cc1cc2* GFP-CC1 and *cc1cc2* GFP-CC1ΔN120 lines and A. Molina, G. Bissoli and E. López-Solanilla for constructive discussions. Live cell imaging was performed with equipment maintained by the Center for Microscopy and Image Analysis (University of Zurich) and Scientific Center for Optical and Electron Microscopy (ScopeM, ETH Zurich). The authors would like to thank Lotte Bald and Rainer Waadt (COS Heidelberg) for cloning pUB10:SYP122-pHusion and pGGC-pHGFP, respectively. We thank David Vukovic (Plückthun group/University of Zurich) for advice on imaging and data analysis. The research leading to these results has received funding from the Peter und Traudl Engelhorn-Stiftung to C.K., Swiss National Foundation to C.S.-R. (SNF 2-77212-15; A.M.), ETHZ foundation to C.S.-R. (0-20172-16; H-Y.H.), Heinz Imhof to C.S.-R. (2-72160-16; A.I.H.), Grants-in-Aid for Scientific Research from the Ministry of Education, Culture, Sports, Science, and Technology, Japan to T.K. (15H05956), Innovational Research Incentives Scheme Vici of The Netherlands Organisation for Scientific Research (NWO) and the Horizon programme of the Netherlands Genomics Initiative through grants to M.R., German Research Foundation (Deutsche Forschungsgemeinschaft, DFG) to K.S. (SFB 1101-TPA02).

## Article and author information

**Author details**

**Christopher Kesten**

Department of Biology, ETH Zurich, 8092 Zurich, Switzerland

Contribution: Conception and design, acquisition, analysis and interpretation of data, drafting and revising the article.

Competing interests: The authors declare no competing interests.

**Francisco M. Gámez-Arjona**

Department of Biology, ETH Zurich, 8092 Zurich, Switzerland

Contribution: Conception and design, acquisition, analysis and interpretation of data, drafting and revising the article.

Competing interests: The authors declare no competing interests.

**Stefan Scholl**

Centre for Organismal Studies, Plant Developmental Biology, Heidelberg University, 69120 Heidelberg, Germany

Contribution: Contributed unpublished essential material.

Competing interests: The authors declare no competing interests.

**Alexandra Menna**

Department of Biology, ETH Zurich, 8092 Zurich, Switzerland

Contribution: Acquisition, analysis, and interpretation of data.

Competing interests: The authors declare no competing interests.

**Susanne Dora**

Department of Biology, ETH Zurich, 8092 Zurich, Switzerland

Contribution: Acquisition, analysis, and interpretation of data.

Competing interests: The authors declare no competing interests.

**Apolonio Ignacio Huerta**

Department of Biology, ETH Zurich, 8092 Zurich, Switzerland

Contribution: Experimental design.

Competing interests: The authors declare no competing interests.

**Hsin-Yao Huang**

Department of Biology, ETH Zurich, 8092 Zurich, Switzerland

Contribution: Acquisition of data.

Competing interests: The authors declare no competing interests.

**Nico Tintor**

Department of Phytopathology, University of Amsterdam, Amsterdam, The Netherlands

Contribution: Contributed unpublished essential material.

Competing interests: The authors declare no competing interests.

**Martjin Rep**

Department of Phytopathology, University of Amsterdam, Amsterdam, The Netherlands

Contribution: Contributed unpublished essential material.

Competing interests: The authors declare no competing interests.

**Toshinori Kinoshita**

Institute of Transformative Bio-Molecules (WPI-ITbM), Nagoya University, Chikusa, Nagoya, Japan Division of Biological Science, Graduate School of Science, Nagoya University, Chikusa, Nagoya, Japan

Contribution: Contributed essential material.

Competing interests: The authors declare no competing interests.

**Melanie Krebs**

Centre for Organismal Studies, Cell Biology, Heidelberg University, 69120 Heidelberg, Germany

Contribution: Analysis and interpretation of data, contribution of unpublished essential material, revising the article.

Competing interests: The authors declare no competing interests.

**Karin Schumacher**

Centre for Organismal Studies, Cell Biology, Heidelberg University, 69120 Heidelberg, Germany

Contribution: Conception and design, revising the article.

Competing interests: The authors declare no competing interests.

**Clara Sánchez-Rodríguez**

Department of Biology, ETH Zurich, 8092 Zurich, Switzerland

Contribution: Conception and design, analysis and interpretation of data, drafting and revising the article.

Competing interests: The authors declare no competing interests.

## Supplementary Movie Legends

Supplementary Movie 1: Fo5176 hyphae cause immediate depletion of cellulose synthase complexes and cortical microtubules from the plasma membrane and cell cortex.

Supplementary Movie 2: Fo5176 elicitors cause immediate depletion of cellulose synthase complexes and cortical microtubules from the plasma membrane and cell cortex.

Supplementary Movie 3: Fo5176 elicitors cause immediate root growth reduction.

Supplementary Movie 4: Depletion of cellulose synthase complexes and cortical microtubules can be attenuated by buffering the media with MES.

Supplementary Movie 5: Fo5176 elicitors cause immediate acidification of the apoplastic pH.

---

**Supplementary Figure 1:**
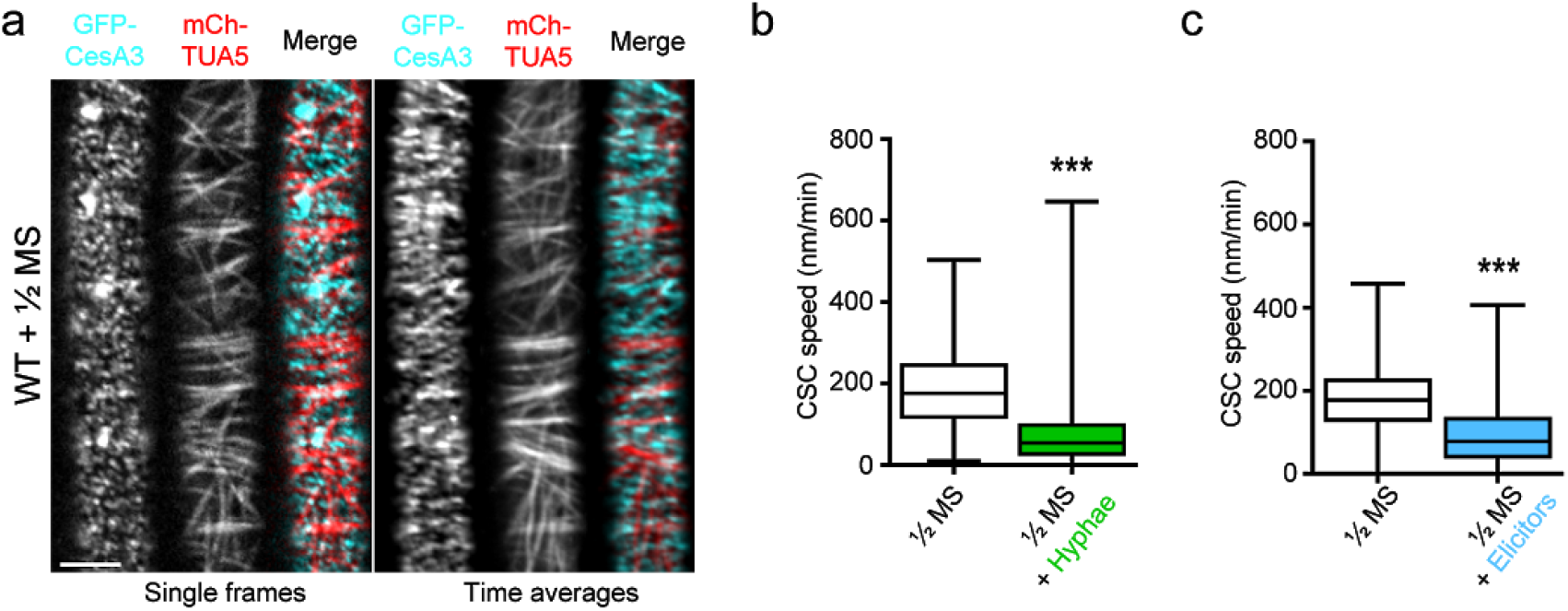
Fo5176 hyphae and elicitors cause immediate slow down of PM-localized cellulose synthase complexes. **a** Representative image of 5 day-old wild-type (WT) GFP-CesA3 and mCh-TUA5 dual-labeled root epidermal cell upon 5 min of half MS treatment (left panels; single frame, right panels; time average projections). Scale bar = 5 μm. **b.** Quantification of GFP-CesA3 speed at the plasma membrane of WT root cells 5 min after half MS treatment (as depicted in **a**) or Fo5176 contact (as depicted in **Fig. 1a**). Box plots: center lines show the medians; box limits indicate the 25th and 75th percentiles; whiskers extend to the minimum and maximum. N ≥ 369 particles from 6 cells and 6 roots and 3 independent experiments; Welch’s unpaired *t*-test; *** p-value ≤ 0.001. **c** Quantification of GFP-CesA3 speed at the plasma membrane of WT root cells 5 min after half MS or half MS + elicitors treatment (as depicted in **Fig. 1f**). Box plots as described in **b**. N ≥ 647 particles from 9 cells and 6 roots and 3 independent experiments; Welch’s unpaired *t*-test; *** p-value ≤ 0.001.

**Supplementary Figure 2:**
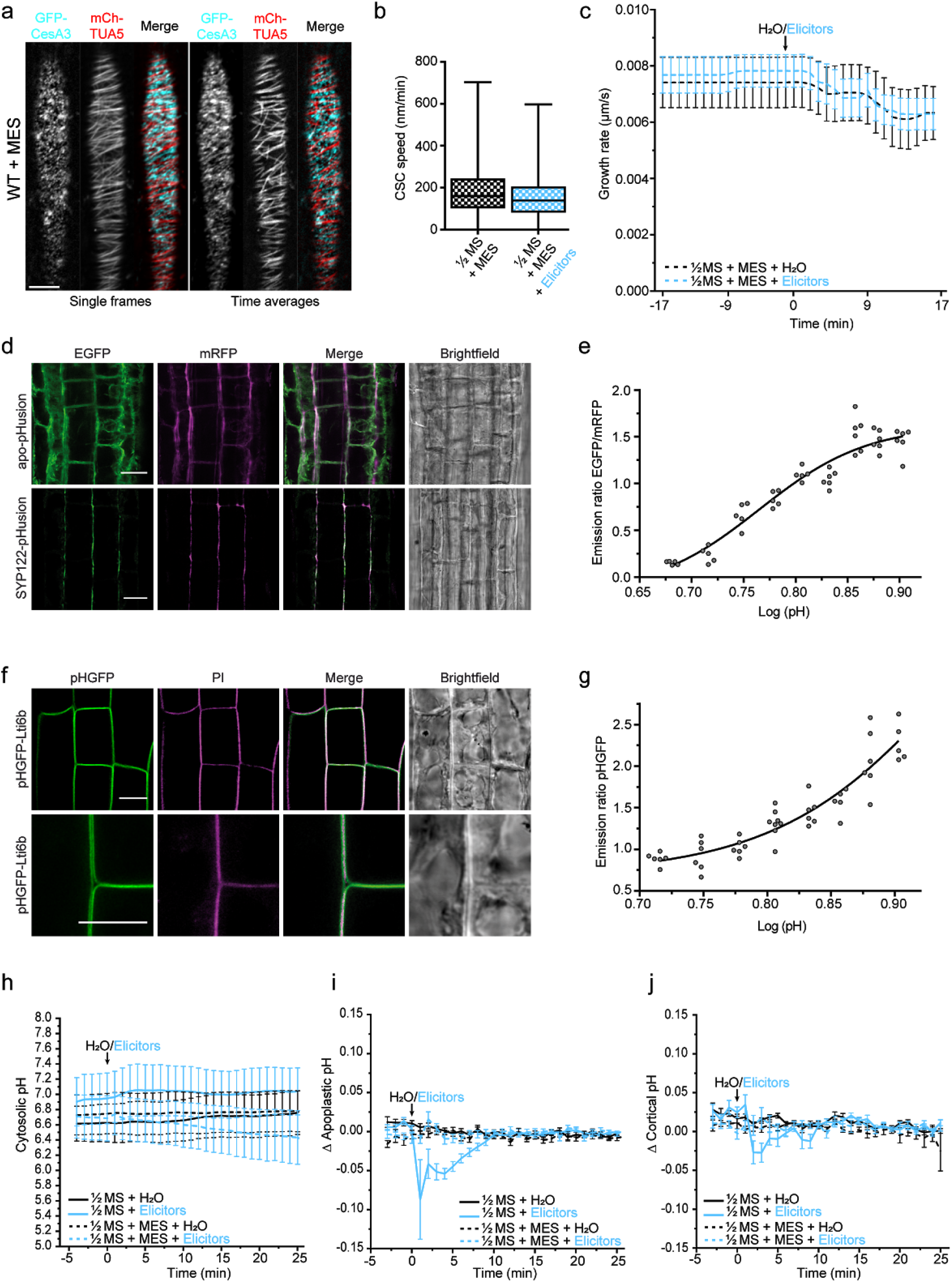
Fo5176-induced CSC-MT alterations and immediate acidification of the PM interface, measured with newly designed sensors, can be buffered with MES. **a** Representative image of a 5 day-old wild-type (WT) GFP-CesA3 and mCh-TUA5 dual-labeled root epidermal cell upon 5 min of half MS + 5 mM MES (left panels; single frame, right panels; time average projections). Scale bar = 5 μm. **b** Quantification of GFP-CesA3 speed at the plasma membrane of WT root cells 5 min after being exposed to half MS + 5 mM MES and half MS + 5 mM MES + Elicitors treatment as depicted in **a** and **Fig. 2a**, respectively. Box plots: center lines show the medians; box limits indicate the 25th and 75th percentiles; whiskers extend to the minimum and maximum. N ≥ 786 particles from 13 cells and 11 roots and 3 independent experiments. **c** Growth rate of roots exposed to fungal elicitors, analyzed from images as in **a**. After 17 min of growth in half MS, 5 mM MES or elicitors + 5 mM MES were added and the growth rate was measured for an additional 17 min. Average growth rate before treatment (−17 to 0 min): 5 mM MES: 0.0074 ± 0.0009 μm/s; Elicitors + 5 mM MES: 0.0068 ± 0.0010 μm/s. Average growth rate after treatment (0-17 min): 5 mM MES: 0.0077 ± 0.0024 μm/s; Elicitors + 5 mM MES: 0.0069 ± 0.0020 μm/s. Values are mean +/-SEM, N ≥ 11 seedlings from 3 independent experiments. Welch’s unpaired *t*-test for roots before and after elicitors treatment; p-value = 0.33. **d** Representative images of 6 days-old wild-type root epidermal and cortex cells expressing apo-pHusion or SYP122-phusion. Remarkably, much less intracellular signal of pHusion is observed for SYP122-pHusion than for apo-pHusion. Scale bar = 20 µm. **e** *In vivo* calibration of SYP122-pHusion in 6 day-old roots. Dots represent individual samples with N ≥ 5 seedlings per standard buffer. Data points were fitted using sigmoidal regression. **f** Representative images of 6 days-old wild-type root epidermal and cortex cells expressing pHGFP-Lti6b. Cell walls of seedlings were counterstained with 3 µg/ml propidium iodide to illustrate plasma membrane localization of pHGFP-Lti6b of two adjacent cells. Scale bars = 10 µm. **g** *In vivo* calibration of pHGFP-Lti6b in 6 day-old roots. Dots represent individual samples with N ≥ 5 seedlings per standard buffer. Data points were fitted using sigmoidal regression. **h** Apoplastic pH variation of WT roots expressing the pH_cyto_ free pHGFP sensor over time, either in half MS or half MS + 5 mM MES. Imaging started 5 min before either H_2_O or a fungal elicitor mix was added (0 min). Values are mean +/-SEM, N ≥ 12 seedlings from 3 independent experiments. RM two-way ANOVA on half MS + H_2_O vs half MS + elicitors: p = 0.42 (treatment), p = 0.06 (time), p = 0.08 (treatment x time). **i** Apoplastic ∆pH of WT roots expressing the pH_apo_ SYP122-pHusion sensor over time, either in half MS or half MS + 5 mM MES. The pH was calculated from image series as in **Fig. 2d**. Imaging started 5 min before either H_2_O or a fungal elicitors were added (0 min). Values are mean +/-SEM, N ≥ 16 seedlings from 3 independent experiments. RM two-way ANOVA on half MS + H_2_O vs half MS + elicitors: p ≤ 0.001 (treatment), p ≤ 0.001 (time), p ≤ 0.001 (treatment x time). **j** Cortical ∆pH of WT roots expressing the pH_cortical_ pHGFP-Lti6b sensor over time, either in half MS or half MS + 5 mM MES. Imaging started 5 min before either H_2_O or a fungal elicitor mix was added (0 min). Mixed effects model on half MS + H_2_O vs half MS + elicitors: p ≤ 0.05 (treatment), p ≤ 0.001 (time), p ≤ 0.001 (treatment x time).

**Supplementary Figure 3:**
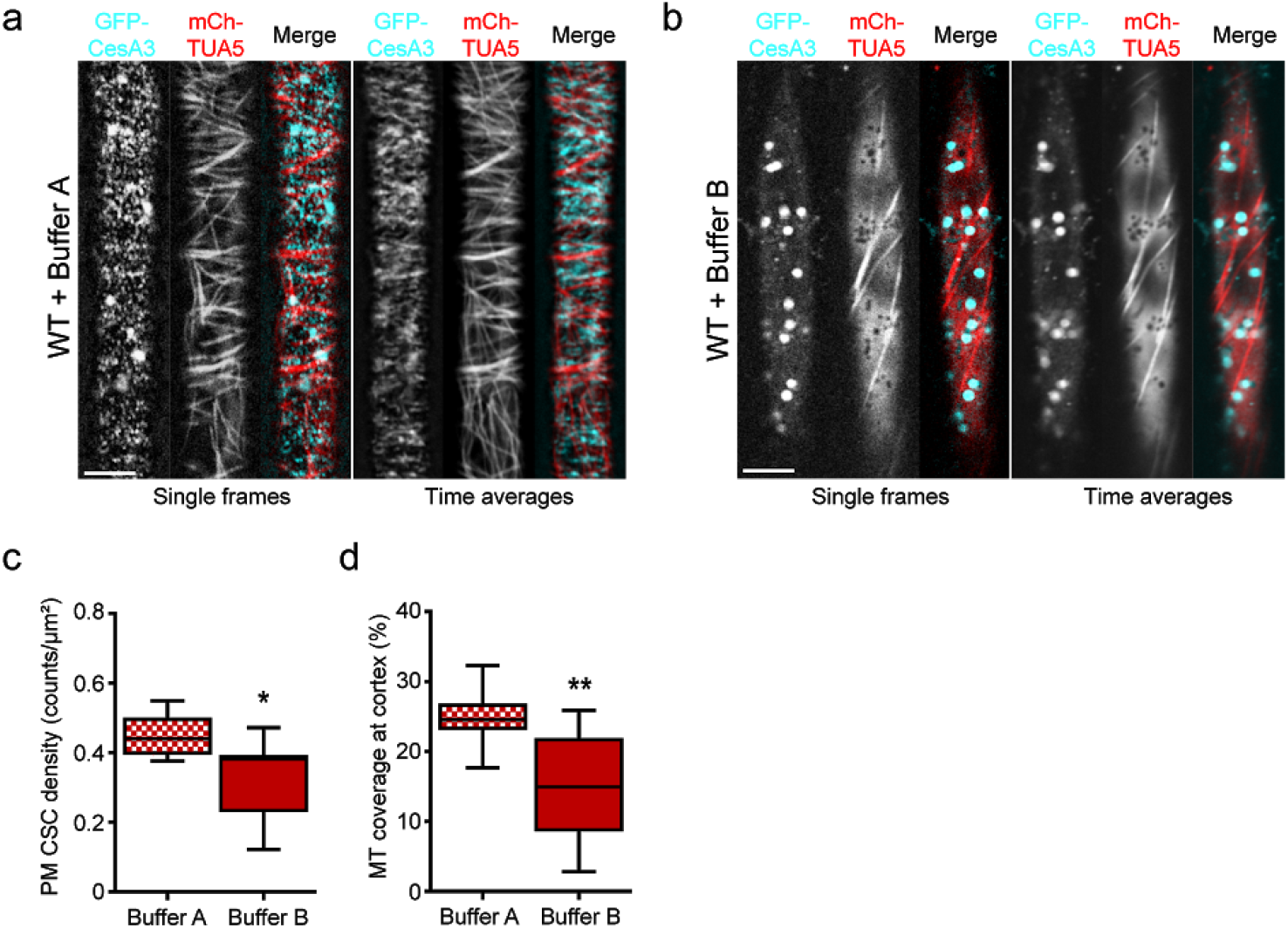
Effect of pH changes at the PM interface on the cellulose synthesis machinery. **a** Representative image of a 5 day-old wild-type (WT) GFP-CesA3 and mCh-TUA5 dual-labeled root epidermal cell upon 5 min treatment with Buffer A (25 mM MES-BTP pH 5.4, 25 mM CH_3_COONH_4_) (left panels; single frame, right panels; time average projections). Scale bar = 5 μm. **b** Representative image of a 5 WT GFP-CesA3 and mCh-TUA5 dual-labeled root epidermal cell upon 5 min treatment with Buffer B (25 mM MES-BTP pH 5.4) (left panels; single frame, right panels; time average projections). Scale bar = 5 μm. **c** Quantification of GFP-CesA3 density at the plasma membrane of WT root cells 5 min after exposure to Buffer A and B as shown in **a** and **b**, respectively. Box plots: center lines show the medians; box limits indicate the 25th and 75th percentiles; whiskers extend to the minimum and maximum. N ≥ 7 cells from 4 roots and 3 independent experiments; Welch’s unpaired *t*-test; * p-value ≤ 0.05. **d** Quantification of microtubule density at cell cortex of WT root cells 5 min after exposure to Buffer A and B as shown in **a** and **b**, respectively. Box plots as described in **c**. N ≥ 7 cells from 4 roots and 3 independent experiments; Welch’s unpaired *t*-test; ** p-value ≤ 0.01.

**Supplementary Figure 4:**
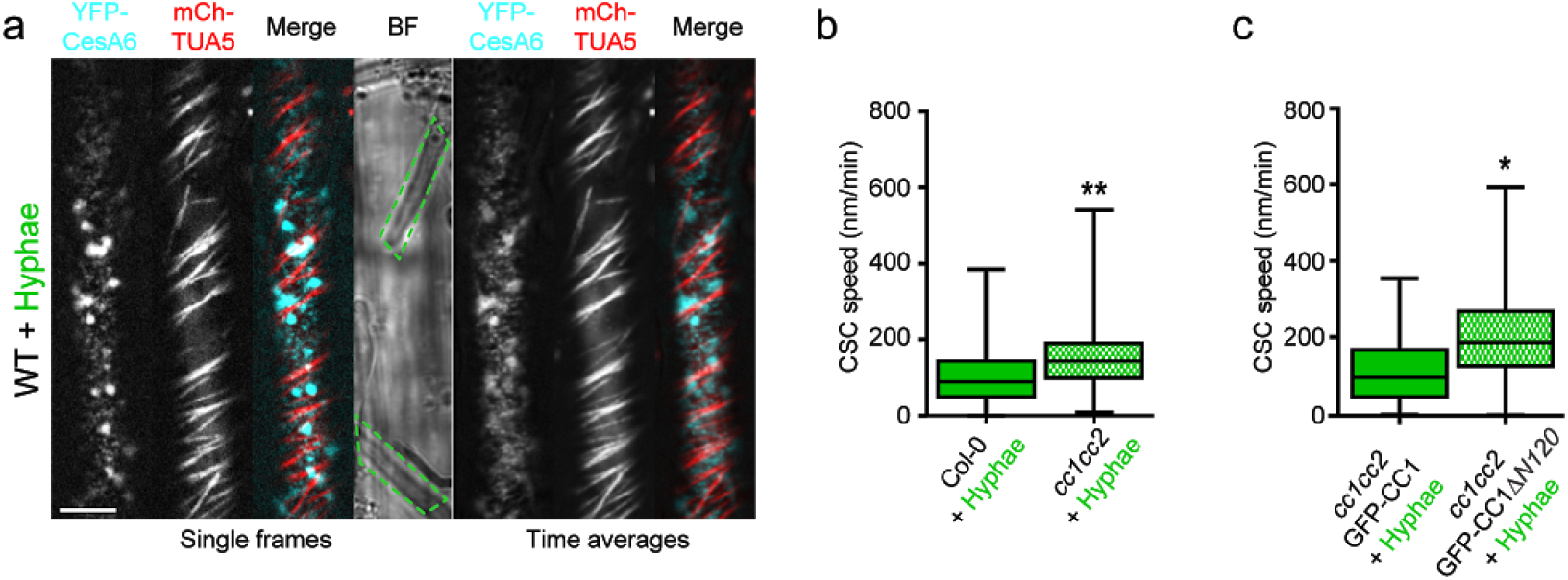
Fo5176 hyphae do not alter the speed of PM-localized cellulose synthase complexes in *cc1cc2* or *cc1cc2* GFP-CC1ΔN120 complemented lines. **a** Representative image of 5 day-old wild-type (WT) YFP-CesA6 and mCh-TUA5 dual-labeled root epidermal cells upon Fo5176 hyphae contact (left panels; single frame, right panels; time average projections). Fo5176 hypha is defined by a green dashed line. Scale bar = 5 μm. **b** Quantification of YFP-CesA6 speed at the plasma membrane of WT WT root cells, as depicted in **a,** and *cc1cc2* background, as depicted in **Fig. 3a,** upon Fo5176 hyphae contact. Box plots: center lines show the medians; box limits indicate the 25th and 75th percentiles; whiskers extend to the minimum and maximum. N ≥ 368 particles from 13 cells and 10 roots and 3 independent experiments; Welch’s unpaired *t*-test; ** p-value ≤ 0.01. **c** Quantification of GFP-CC1 and GFP-CC1ΔN120 speed at the plasma membrane of *cc1cc2* root cells upon Fo5176 hyphae contact (as depicted in **Fig. 3e and f**). Box plots as described in **b**. N ≥ 288 particles from 12 cells and 9 roots and 3 independent experiments; Welch’s unpaired *t*-test; * p-value ≤ 0.05.

**Supplementary Figure 5:**
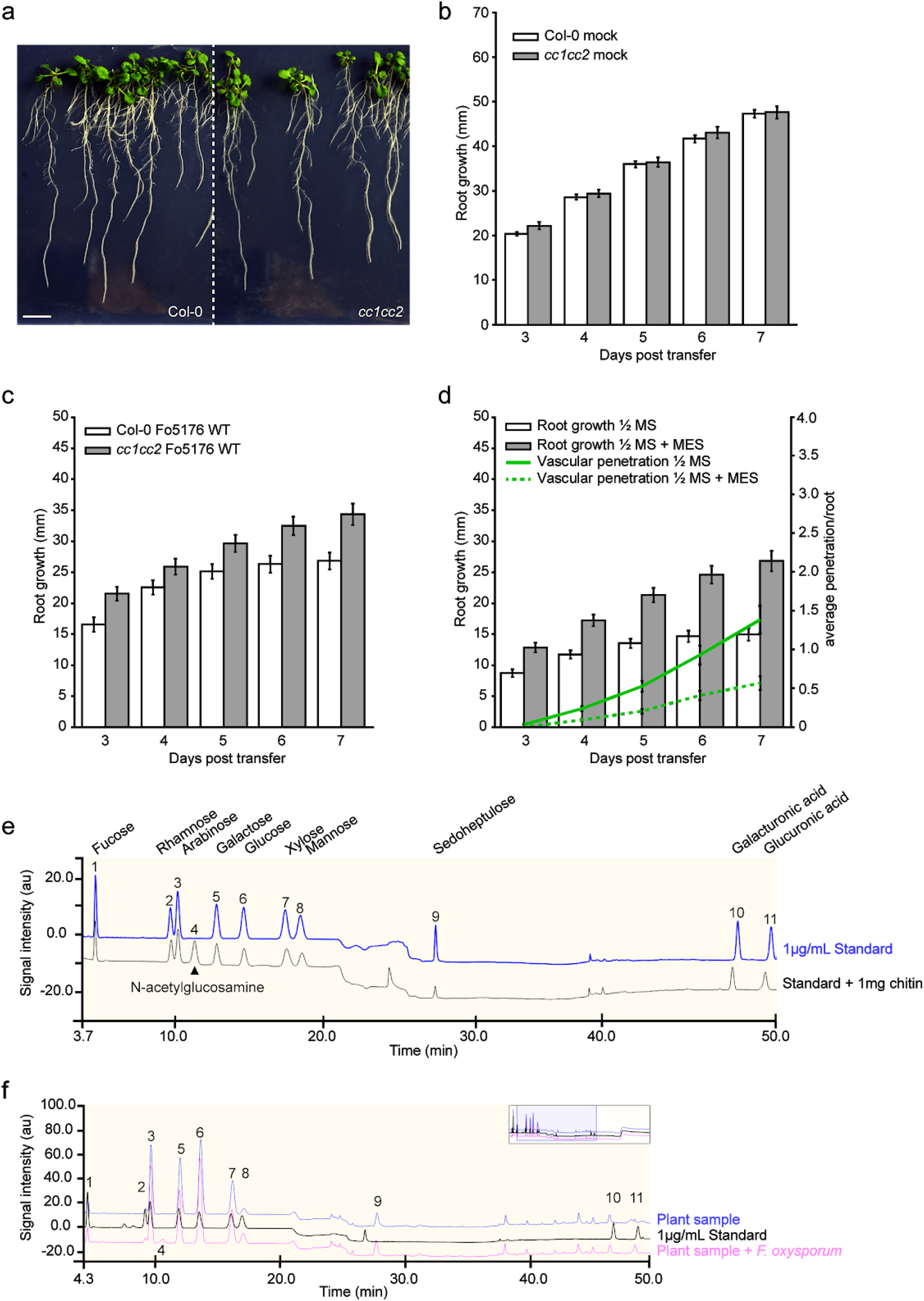
*cc1cc2* resistance to Fo5176 root colonization can be mimicked in wild-type plants by buffering the growth media. **a** Representative image of wild-type (WT, Col-0) and *cc1cc2* plants 7 days post transfer to half MS control plates. Scale bar = 10 mm. **b** Quantification of root elongation of plants grown as in **a**. Values are mean +/-SEM, N ≥ 103 plants from 4 independent experiments. RM two-way ANOVA: p = 0.6506 (genotype), p ≤ 0.001 (time), p = 0.3310 (genotype x time). **c** Quantification of root elongation of plants 7 days after inoculation with Fo5176 WT spores. Values are mean +/-SEM, N ≥ 33 plants from 3 independent experiments. RM two-way ANOVA: p ≤ 0.01 (genotype), p ≤ 0.001 (time), p ≤ 0.05 (genotype x time). **d** Quantification of root elongation and vascular penetration of WT plants at various days post inoculation with Fo5176 pSIX1::GFP w/o 5 mM MES. Values are mean +/-SEM, N ≥ 33 plants from 3 independent experiments. RM two-way ANOVA on root growth: p ≤ 0.001 (genotype), p ≤ 0.001 (time), p ≤ 0.001 (genotype x time). RM two-way ANOVA on vascular penetration rate: p ≤ 0.001 (genotype), p ≤ 0.001 (time), p ≤ 0.001 (genotype x time). **e** Monosaccharide elution profile of a 1μg/mL standard (blue) compared to a 1:5 dilution of a 1μg/mL standard with 1mg of chitin subjected to both the Saeman and weak hydrolyses. N-acetylglucosamine (arrow head, peak 4) derived from chitin elutes in between the arabinose and galactose peaks without interference. **f** Monosaccharide elution profile of a 1μg/mL standard (black) compared to 1:10 dilutions of hydrolyzed samples derived roots after 12 days of either mock (blue) or Fo5176 treatment (pink). The N-acetylglucosamine peak (4) is only identifiable in hydrolyzed infected samples (pink).

**Supplementary Figure 6:**
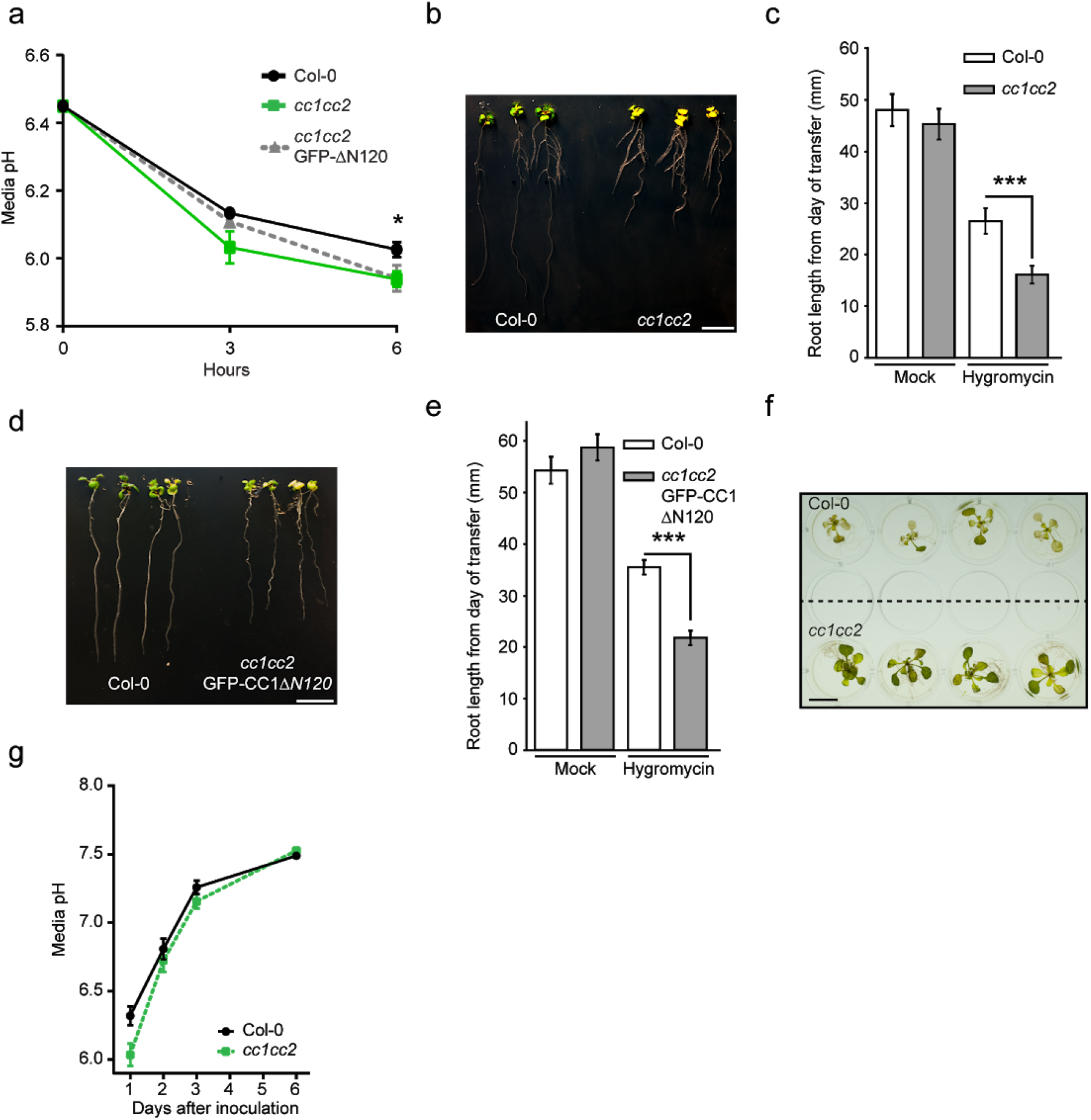
*cc1cc2* mutants rapidly acidify the media, are hypersensitive to hygromycin B and show a delay on Fo5176-induced media alkalinization. **a** Media pH development over time when 10 day-old wild-type (WT, Col-0), *cc1cc2,* and *cc1cc2* GFP-CC1ΔN120 plants were transferred to a liquid, alkaline media (pH 6.45). Values are mean +/-SEM, N = 24 plants from 3 independent experiments. Welch’s unpaired *t*-test; * p-value ≤ 0.05. **b** Representative image of WT and *cc1cc2* plants 7 days after being transferred to half MS + 5 µg/ml hygromycin B plates. Scale bar = 10 mm. **c** Quantification of root elongation of plants grown as in **b**. Values are mean +/-SEM, N ≥ 15 plants from 3 independent experiments. Welch’s unpaired *t*-test; *** p-value ≤ 0.001. **d** Representative image of WT and *cc1cc2* GFP-CC1ΔN120 plants 7 days after being transferred half MS + 5 µg/ml hygromycin B plates. Scale bar = 10 mm. **e** Quantification of root elongation of plants grown as in **d**. Values are mean +/-SEM, N ≥ 16 plants from 3 independent experiments. Welch’s unpaired *t*-test; *** p-value ≤ 0.001. **f** Representative image of 10 day-old WT and *cc1cc2* plants, which were inoculated with Fo5176 pSIX1:GFP spores and grown for an additional 7 days in liquid half MS media. Scale bar = 10 mm. **g** Media pH development over time when 10 day-old WT or *cc1cc2* plants were inoculated with Fo5176 pSIX1:GFP spores as in **f**. Values are mean +/-SEM, N = 36 plants from 3 independent experiments. RM two-way ANOVA: p = 0.069 (genotype), p ≤ 0.001 (time), p ≤ 0.05 (genotype x time).

**Supplementary Table 1:**
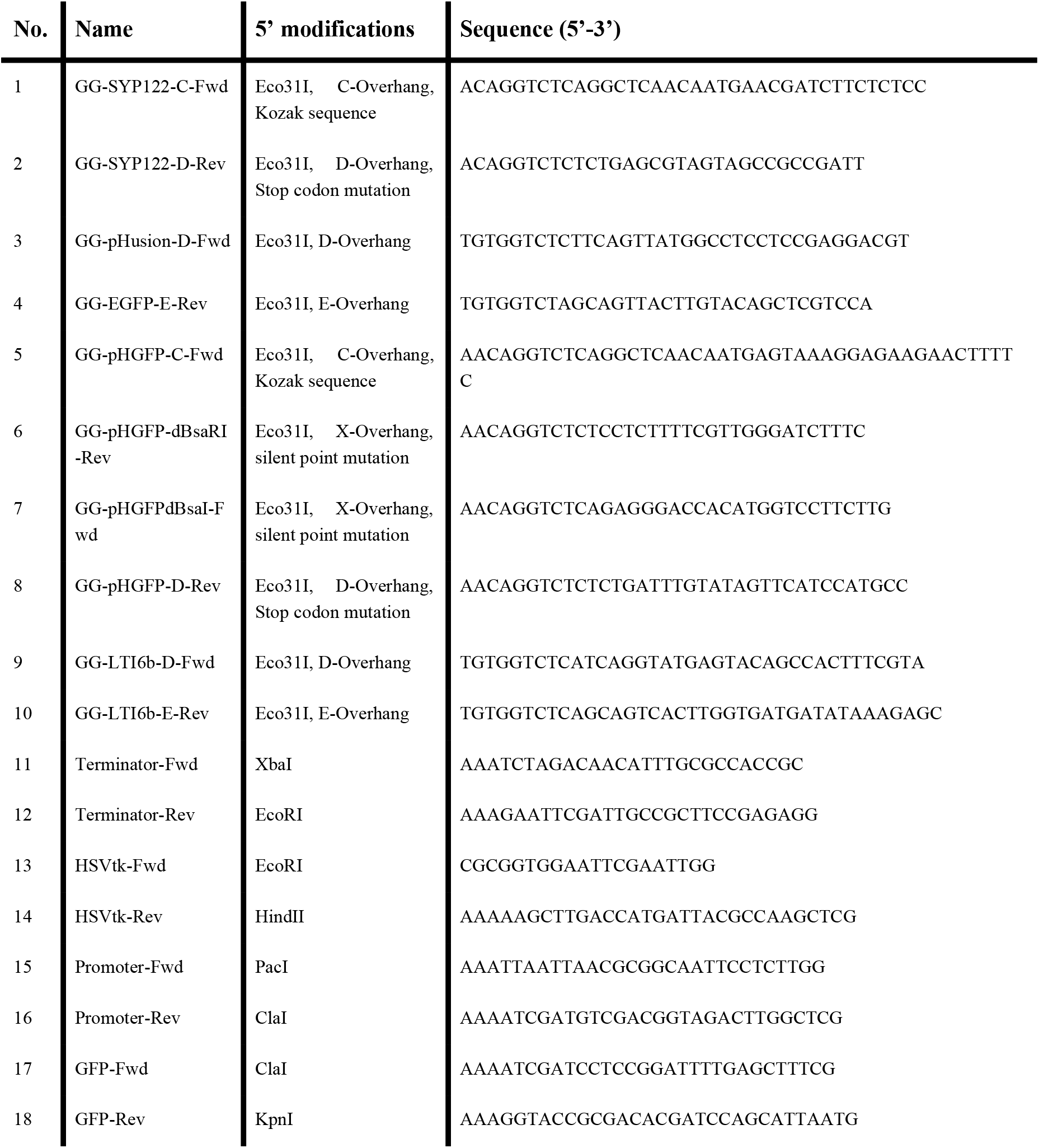
Primers used in the manuscript (5’-3’).

## References

Assaad FF, Qiu J-L, Youngs H, Ehrhardt D, Zimmerli L, Kalde M, Wanner G, Peck SC, Edwards H, Ramonell K, Somerville CR & Thordal-Christensen H (2004) The PEN1 syntaxin defines a novel cellular compartment upon fungal attack and is required for the timely assembly of papillae. Mol. Biol. Cell 15: 5118–5129

Baldrich P, Kakar K, Siré C, Moreno AB, Berger A, García-Chapa M, López-Moya JJ, Riechmann JL & San Segundo B (2014) Small RNA profiling reveals regulation of Arabidopsis miR168 and heterochromatic siRNA415 in response to fungal elicitors. BMC Genomics 15: 1083

Barbez E, Dünser K, Gaidora A, Lendl T & Busch W (2017) Auxin steers root cell expansion via apoplastic pH regulation in Arabidopsis thaliana. Proc. Natl. Acad. Sci. U. S. A. 114: E4884–E4893

Behera S, Xu Z, Luoni L, Bonza C, Doccula FG, DeMichelis MI, Morris RJ, Schwarzländer M & Costa A (2018) Cellular Ca2+ signals generate defined pH signatures in plants. Plant Cell Available at: http://dx.doi.org/10.1105/tpc.18.00655

Benschop JJ, Mohammed S, O’Flaherty M, Heck AJR, Slijper M & Menke FLH (2007) Quantitative phosphoproteomics of early elicitor signaling in Arabidopsis. Mol. Cell. Proteomics 6: 1198–1214

Bugbee BG & Salisbury FB (1985) An evaluation of MES (2(N-Morpholino)ethanesulfonic acid) and amberlite IRC-50 as pH buffers for nutrient solution studies. J. Plant Nutr. 8: 567–583

Cosgrove DJ (2018) Nanoscale structure, mechanics and growth of epidermal cell walls. Curr. Opin. Plant Biol. 46: 77–86

Cutler SR, Ehrhardt DW, Griffitts JS & Somerville CR (2000) Random GFP∷cDNA fusions enable visualization of subcellular structures in cells of Arabidopsis at a high frequency. Proc. Natl. Acad. Sci. U. S. A. 97: 3718–3723

Desprez T, Juraniec M, Crowell EF, Jouy H, Pochylova Z, Parcy F, Höfte H, Gonneau M & Vernhettes S (2007) Organization of cellulose synthase complexes involved in primary cell wall synthesis in Arabidopsis thaliana. Proc. Natl. Acad. Sci. U. S. A. 104: 15572–15577

Di Pietro A, Garcia-Maceira FI, Meglecz E & Roncero MIG (2001) A MAP kinase of the vascular wilt fungus Fusarium oxysporum is essential for root penetration and pathogenesis. Mol. Microbiol. 39: 1140–1152

van der Does HC, Duyvesteijn RGE, Goltstein PM, van Schie CCN, Manders EMM, Cornelissen BJC & Rep M (2008) Expression of effector gene SIX1 of Fusarium oxysporum requires living plant cells. Fungal Genet. Biol. 45: 1257–1264

Elmore JM & Coaker G (2011) The role of the plasma membrane H+-ATPase in plant-microbe interactions. Mol. Plant 4: 416–427

Endler A, Kesten C, Schneider R, Zhang Y, Ivakov A, Froehlich A, Funke N & Persson S (2015) A Mechanism for Sustained Cellulose Synthesis during Salt Stress. Cell 162: 1353–1364

Falhof J, Pedersen JT, Fuglsang AT & Palmgren M (2016) Plasma Membrane H(+)-ATPase Regulation in the Center of Plant Physiology. Mol. Plant 9: 323–337

Felle HH (2001) pH: signal and messenger in plant cells. Plant Biol. 3: 577–591

Felle HH, Herrmann A, Hückelhoven R & Kogel K-H (2005) Root-to-shoot signalling: apoplastic alkalinization, a general stress response and defence factor in barley (Hordeum vulgare). Protoplasma 227: 17–24

Felle HH, Waller F, Molitor A & Kogel K-H (2009) The Mycorrhiza Fungus Piriformospora indica Induces Fast Root-Surface pH Signaling and Primes Systemic Alkalinization of the Leaf Apoplast Upon Powdery Mildew Infection. Mol. Plant. Microbe. Interact. 22: 1179–1185

Fendrych M, Leung J & Friml J (2016) TIR1/AFB-Aux/IAA auxin perception mediates rapid cell wall acidification and growth of Arabidopsis hypocotyls. Elife 5: Available at: http://dx.doi.org/10.7554/eLife.19048

Fendrych M, Van Hautegem T, Van Durme M, Olvera-Carrillo Y, Huysmans M, Karimi M, Lippens S, Guérin CJ, Krebs M, Schumacher K & Nowack MK (2014) Programmed cell death controlled by ANAC033/SOMBRERO determines root cap organ size in Arabidopsis. Curr. Biol. 24: 931–940

Feng W, Lindner H, Robbins NE 2nd & Dinneny JR (2016) Growing Out of Stress: The Role of Cell- and Organ-Scale Growth Control in Plant Water-Stress Responses. Plant Cell 28: 1769–1782

Gao D, Knight MR, Trewavas AJ, Sattelmacher B & Plieth C (2004) Self-Reporting Arabidopsis Expressing pH and [Ca2+] Indicators Unveil Ion Dynamics in the Cytoplasm and in the Apoplast under Abiotic Stress. Plant Physiol. 134: 898–908

Geilfus C-M (2017) The pH of the Apoplast: Dynamic Factor with Functional Impact Under Stress. Mol. Plant 10: 1371–1386

Gjetting KSK, Ytting CK, Schulz A & Fuglsang AT (2012) Live imaging of intra- and extracellular pH in plants using pHusion, a novel genetically encoded biosensor. J. Exp. Bot. 63: 3207–3218

Good NE, Winget GD, Winter W, Connolly TN, Izawa S & Singh RMM (1966) Hydrogen Ion Buffers for Biological Research*. Biochemistry 5: 467–477

Gutierrez R, Lindeboom JJ, Paredez AR, Emons AMC & Ehrhardt DW (2009) Arabidopsis cortical microtubules position cellulose synthase delivery to the plasma membrane and interact with cellulose synthase trafficking compartments. Nat. Cell Biol. 11: 797–806

Hardham AR, Takemoto D & White RG (2008) Rapid and dynamic subcellular reorganization following mechanical stimulation of Arabidopsis epidermal cells mimics responses to fungal and oomycete attack. BMC Plant Biol. 8: 63

Haruta M, Burch HL, Nelson RB, Barrett-Wilt G, Kline KG, Mohsin SB, Young JC, Otegui MS & Sussman MR (2010) Molecular characterization of mutant Arabidopsis plants with reduced plasma membrane proton pump activity. J. Biol. Chem. 285: 17918–17929

Haruta M, Gray WM & Sussman MR (2015) Regulation of the plasma membrane proton pump (H(+)-ATPase) by phosphorylation. Curr. Opin. Plant Biol. 28: 68–75

Hayashi Y, Nakamura S, Takemiya A, Takahashi Y, Shimazaki K-I & Kinoshita T (2010) Biochemical Characterization of In Vitro Phosphorylation and Dephosphorylation of the Plasma Membrane H-ATPase. Plant Cell Physiol. 51: 1186–1196

Heiple JM & Taylor DL (1980) Intracellular pH in single motile cells. J. Cell Biol.86: 885–890

Hellens RP, Edwards EA, Leyland NR, Bean S & Mullineaux PM (2000) pGreen: a versatile and flexible binary Ti vector for Agrobacterium-mediated plant transformation. Plant Mol. Biol. 42: 819–832

Hostettler C, Kölling K, Santelia D, Streb S, Kötting O & Zeeman SC (2011) Analysis of starch metabolism in chloroplasts. Methods Mol. Biol. 775: 387–410

Houterman PM, Cornelissen BJC & Rep M (2008) Suppression of plant resistance gene-based immunity by a fungal effector. PLoS Pathog. 4: e1000061

Houterman PM, Speijer D, Dekker HL, DE Koster CG, Cornelissen BJC & Rep M (2007) The mixed xylem sap proteome of Fusarium oxysporum-infected tomato plants. Mol. Plant Pathol. 8: 215–221

Inoue S-I & Kinoshita T (2017) Blue Light Regulation of Stomatal Opening and the Plasma Membrane H+-ATPase. Plant Physiol. 174: 531–538

Jeworutzki E, Roelfsema MRG, Anschütz U, Krol E, Elzenga JTM, Felix G, Boller T, Hedrich R & Becker D (2010) Early signaling through the Arabidopsis pattern recognition receptors FLS2 and EFR involves Ca-associated opening of plasma membrane anion channels. Plant J. 62: 367–378

Kesten C, Menna A & Sánchez-Rodríguez C (2017) Regulation of cellulose synthesis in response to stress. Curr. Opin. Plant Biol. 40: 106–113

Kesten C, Schneider R & Persson S (2016) In vitro Microtubule Binding Assay and Dissociation Constant Estimation. BIO-PROTOCOL 6: e1759

Kesten C, Wallmann A, Schneider R, McFarlane HE, Diehl A, Khan GA, van Rossum B-J, Lampugnani ER, Cremer N, Schmieder P, Ford KL, Seiter F, Heazlewood JL, Sanchez-Rodriguez C, Oschkinat H & Persson S (2018) A molecular mechanism for salt stress-induced microtubule array formation in Arabidopsis. bioRxiv: 431031 Available at: https://www.biorxiv.org/content/early/2018/10/11/431031 [Accessed December 14, 2018]

Khang CH, Park S-Y, Lee Y-H & Kang S (2005) A dual selection based, targeted gene replacement tool for Magnaporthe grisea and Fusarium oxysporum. Fungal Genet. Biol. 42: 483–492

Krebs M, Held K, Binder A, Hashimoto K, Den Herder G, Parniske M, Kudla J & Schumacher K (2012) FRET-based genetically encoded sensors allow high-resolution live cell imaging of Ca^2+^ dynamics. Plant J. 69: 181–192

Krebs M & Schumacher K (2013) Live cell imaging of cytoplasmic and nuclear Ca2+ dynamics in Arabidopsis roots. Cold Spring Harb. Protoc. 2013 776–780

Laemmli UK (1970) Cleavage of structural proteins during the assembly of the head of bacteriophage T4. Nature 227: 680–685

Lampropoulos A, Sutikovic Z, Wenzl C, Maegele I, Lohmann JU & Forner J (2013) GreenGate---a novel, versatile, and efficient cloning system for plant transgenesis. PLoS One 8: e83043

Li B, Wang W, Zong Y, Qin G & Tian S (2012) Exploring pathogenic mechanisms of Botrytis cinerea secretome under different ambient pH based on comparative proteomic analysis. J. Proteome Res. 11: 4249–4260

Lindeboom JJ, Nakamura M, Hibbel A, Shundyak K, Gutierrez R, Ketelaar T, Emons AMC, Mulder BM, Kirik V & Ehrhardt DW (2013) A mechanism for reorientation of cortical microtubule arrays driven by microtubule severing. Science 342: 1245533

Liu Y, Visetsouk M, Mynlieff M, Qin H, Lechtreck KF & Yang P (2017) H+- and Na+-elicited rapid changes of the microtubule cytoskeleton in the biflagellated green alga Chlamydomonas. Elife 6: Available at: http://dx.doi.org/10.7554/eLife.26002

Liu Z, Persson S & Sánchez-Rodríguez C (2015) At the border: the plasma membrane-cell wall continuum. J. Exp. Bot. 66: 1553–1563

López-Díaz C, Rahjoo V, Sulyok M, Ghionna V, Martín-Vicente A, Capilla J, Di Pietro A & López-Berges MS (2018) Fusaric acid contributes to virulence of Fusarium oxysporum on plant and mammalian hosts. Mol. Plant Pathol. 19: 440–453

Luo Y, Scholl S, Doering A, Zhang Y, Irani NG, Di Rubbo S, Neumetzler L, Krishnamoorthy P, Van Houtte I, Mylle E, Bischoff V, Vernhettes S, Winne J, Friml J, Stierhof Y-D, Schumacher K, Persson S & Russinova E (2015) V-ATPase activity in the TGN/EE is required for exocytosis and recycling in Arabidopsis. Nature Plants 1: 15094

Mangano S, Martínez Pacheco J, Marino-Buslje C & Estevez JM (2018) How Does pH Fit in with Oscillating Polar Growth? Trends Plant Sci. 23: 479–489

Martinière A, Gibrat R, Sentenac H, Dumont X, Gaillard I & Paris N (2018) Uncovering pH at both sides of the root plasma membrane interface using noninvasive imaging. Proc. Natl. Acad. Sci. U. S. A. 115: 6488–6493

Masachis S, Segorbe D, Turrà D, Leon-Ruiz M, Fürst U, El Ghalid M, Leonard G, López-Berges MS, Richards TA, Felix G & Di Pietro A (2016) A fungal pathogen secretes plant alkalinizing peptides to increase infection. Nature Microbiology Available at: http://dx.doi.org/10.1038/nmicrobiol.2016.43

McFarlane HE, Döring A & Persson S (2014) The cell biology of cellulose synthesis. Annu. Rev. Plant Biol. 65: 69–94

Model MA (2006) Intensity calibration and shading correction for fluorescence microscopes. Curr. Protoc. Cytom. Chapter 10: Unit10.14

Model MA & Burkhardt JK (2001) A standard for calibration and shading correction of a fluorescence microscope. Cytometry 44: 309–316

Morris EC, Griffiths M, Golebiowska A, Mairhofer S, Burr-Hersey J, Goh T, von Wangenheim D, Atkinson B, Sturrock CJ, Lynch JP, Vissenberg K, Ritz K, Wells DM, Mooney SJ & Bennett MJ (2017) Shaping 3D Root System Architecture. Curr. Biol. 27: R919–R930

Moseyko N & Feldman LJ (2001) Expression of pH-sensitive green fluorescent protein in Arabidopsis thaliana. Plant Cell Environ. 24: 557–563

Nick P (2013) Microtubules, signalling and abiotic stress. Plant J. 75: 309–323

Nühse TS, Bottrill AR, Jones AME & Peck SC (2007) Quantitative phosphoproteomic analysis of plasma membrane proteins reveals regulatory mechanisms of plant innate immune responses. Plant J. 51: 931–940

Olsson A, Svennelid F, Ek B, Sommarin M & Larsson C (1998) A phosphothreonine residue at the C-terminal end of the plasma membrane H+-ATPase is protected by fusicoccin-induced 14-3-3 binding. Plant Physiol. 118: 551–555

Otsu N (1979) A threshold selection method from gray-level histograms. IEEE Trans. Syst. Man Cybern. 9: 62–66

Paredez AR, Somerville C & Ehrhardt D (2006) Visualization of Cellulose Synthase with Microtubules. Science 312: 1491–1495

Pietro AD, Madrid MP, Caracuel Z, Delgado-Jarana J & Roncero MIG (2003) Fusarium oxysporum: exploring the molecular arsenal of a vascular wilt fungus. Mol. Plant Pathol. 4: 315–325

Pittman JK (2012) Multiple Transport Pathways for Mediating Intracellular pH Homeostasis: The Contribution of H(+)/ion Exchangers. Front. Plant Sci. 3: 11

Platre MP, Noack LC, Doumane M, Bayle V, Simon MLA, Maneta-Peyret L, Fouillen L, Stanislas T, Armengot L, Pejchar P, Caillaud M-C, Potocký M, Čopič A, Moreau P & Jaillais Y (2018) A Combinatorial Lipid Code Shapes the Electrostatic Landscape of Plant Endomembranes. Dev. Cell 45: 465–480.e11

Prager-Khoutorsky M, Khoutorsky A & Bourque CW (2014) Unique interweaved microtubule scaffold mediates osmosensory transduction via physical interaction with TRPV1. Neuron 83: 866–878

Reverchon S & Nasser W (2013) Dickeya ecology, environment sensing and regulation of virulence programme. Environ. Microbiol. Rep. 5: 622–636

Roncero MI, Di Pietro A, Ruiz-Roldán MC, Huertas-González MD, Garcia-Maceira FI, Méglecz E, Jiménez A, Caracuel Z, Sancho-Zapatero R, Hera C, Gómez-Gómez E, Ruiz-Rubio M, González-Verdejo CI & Páez MJ (2000) Role of cell wall-degrading enzymes in pathogenicity of Fusarium oxysporum. Rev. Iberoam. Micol. 17: S47–53

Ruhnow F, Zwicker D & Diez S (2011) Tracking single particles and elongated filaments with nanometer precision. Biophys. J. 100: 2820–2828

Sánchez-Rangel D, Hernández-Domínguez E-E, Pérez-Torres C-A, Ortiz-Castro R, Villafán E, Rodríguez-Haas B, Alonso-Sánchez A, López-Buenfil A, Carrillo-Ortiz N, Hernández-Ramos L & Ibarra-Laclette E (2018) Environmental pH modulates transcriptomic responses in the fungus Fusarium sp. associated with KSHB Euwallacea sp. near fornicatus. BMC Genomics 19: 721

Schindelin J, Arganda-Carreras I, Frise E, Kaynig V, Longair M, Pietzsch T, Preibisch S, Rueden C, Saalfeld S, Schmid B, Tinevez J-Y, White DJ, Hartenstein V, Eliceiri K, Tomancak P & Cardona A (2012) Fiji: an open-source platform for biological-image analysis. Nat. Methods 9: 676–682

Schulte A, Lorenzen I, Böttcher M & Plieth C (2006) A novel fluorescent pH probe for expression in plants. Plant Methods 2: 7

Smakowska E, Kong J, Busch W & Belkhadir Y (2016) Organ-specific regulation of growth-defense tradeoffs by plants. Curr. Opin. Plant Biol. 29: 129–137

Staal M, De Cnodder T, Simon D, Vandenbussche F, Van der Straeten D, Verbelen J-P, Elzenga T & Vissenberg K (2011) Apoplastic alkalinization is instrumental for the inhibition of cell elongation in the Arabidopsis root by the ethylene precursor1-aminocyclopropane-1-carboxylic acid. Plant Physiol. 155: 2049–2055

Stahl RS, Grossl P & Bugbee B (1999) Effect of 2(N-morpholino)ethane)sulfonic acid (MES) on the growth and tissue composition of cucumber. J. Plant Nutr. 22: 315–330

Stecker KE, Minkoff BB & Sussman MR (2014) Phosphoproteomic Analyses Reveal Early Signaling Events in the Osmotic Stress Response. Plant Physiol. 165: 1171–1187

Stegmann M, Monaghan J, Smakowska-Luzan E, Rovenich H, Lehner A, Holton N, Belkhadir Y & Zipfel C (2017) The receptor kinase FER is a RALF-regulated scaffold controlling plant immune signaling. Science 355: 287–289

Sze H & Chanroj S (2018) Plant Endomembrane Dynamics: Studies of K+/H+ Antiporters Provide Insights on the Effects of pH and Ion Homeostasis. Plant Physiol. 177: 875–895

Takken FLW, Van Wijk R, Michielse CB, Houterman PM, Ram AFJ & Cornelissen BJC (2004) A one-step method to convert vectors into binary vectors suited for Agrobacterium-mediated transformation. Curr. Genet. 45: 242–248

Thatcher LF, Gardiner DM, Kazan K & Manners JM (2012) A highly conserved effector in Fusarium oxysporum is required for full virulence on Arabidopsis. Mol. Plant. Microbe. Interact. 25: 180–190

Thevenaz P, Ruttimann UE & Unser M (1998) A pyramid approach to subpixel registration based on intensity. IEEE Trans. Image Process. 7: 27–41

Uemura T, Ueda T, Ohniwa RL, Nakano A, Takeyasu K & Sato MH (2004) Systematic analysis of SNARE molecules in Arabidopsis: dissection of the post-Golgi network in plant cells. Cell Struct. Funct. 29: 49–65

Wang T, McFarlane HE & Persson S (2016) The impact of abiotic factors on cellulose synthesis. J. Exp. Bot. 67: 543–552

Watanabe Y, Meents MJ, McDonnell LM, Barkwill S, Sampathkumar A, Cartwright HN, Demura T, Ehrhardt DW, Samuels AL & Mansfield SD (2015) Visualization of cellulose synthases in Arabidopsis secondary cell walls. Science 350: 198–203

Wolf S (2017) Plant cell wall signalling and receptor-like kinases. Biochem. J 474: 471–492

Yao L-L, Zhou Q, Pei B-L & Li Y-Z (2011) Hydrogen peroxide modulates the dynamic microtubule cytoskeleton during the defence responses to Verticillium dahliae toxins in Arabidopsis. Plant Cell Environ. 34: 1586–1598

Yeats TH, Sorek H, Wemmer DE & Somerville CR (2016a) Cellulose Deficiency Is Enhanced on Hyper Accumulation of Sucrose by a H+-Coupled Sucrose Symporter. Plant Physiol. 171: 110–124

Yeats T, Vellosillo T, Sorek N, Ibáñez AB & Bauer S (2016b) Rapid Determination of Cellulose, Neutral Sugars, and Uronic Acids from Plant Cell Walls by One-step Two-step Hydrolysis and HPAEC-PAD. BIO-PROTOCOL 6: e1978

Yoshida S (1994) Low Temperature-Induced Cytoplasmic Acidosis in Cultured Mung Bean (Vigna radiata [L.] Wilczek) Cells. Plant Physiol. 104: 1131–1138

Zamioudis C, Korteland J, Van Pelt JA, van Hamersveld M, Dombrowski N, Bai Y, Hanson J, Van Verk MC, Ling H-Q, Schulze-Lefert P & Pieterse CMJ (2015) Rhizobacterial volatiles and photosynthesis-related signals coordinate MYB72 expression in Arabidopsis roots during onset of induced systemic resistance and iron-deficiency responses. Plant J. 84: 309–322

